# A combination of expansion microscopy and proximity labelling reveals conserved and unique asymmetric functional hubs at the trypanosome nuclear pore

**DOI:** 10.1101/2024.10.04.616621

**Authors:** Bernardo Papini Gabiatti, Johanna Odenwald, Silke Braune, Timothy Krüger, Martin Zoltner, Susanne Kramer

## Abstract

Nuclear export of mRNAs requires loading the mRNP to the transporter Mex67/Mtr2 in the nucleoplasm, controlled access to the pore by the basket-localized TREX2 complex and mRNA release at the cytoplasmic site by the DEAD-box RNA helicase Dbp5. Asymmetric localisation of nucleoporins (NUPs) and transport components as well as the ATP dependency of Dbp5 ensure unidirectionality of transport. Trypanosomes possess homologues of the mRNA transporter Mex67/Mtr2, but not of TREX2 or Dbp5. Instead, nuclear export is likely fuelled by the GTP/GDP gradient created by the Ran GTPase. However, it remains unclear, how directionality is achieved since the current model of the trypanosomatid pore is mostly symmetric.

We have revisited the architecture of the trypanosome nuclear pore complex using a novel combination of expansion microscopy, proximity labelling and streptavidin imaging. We could confidently assign the NUP76 complex, a known Mex67 interaction platform, to the cytoplasmic site of the pore. The resulting availability of reference proteins for basket, inner ring and cytoplasmic site allowed mapping of all 75 trypanosome proteins with known nuclear pore localisation to a sub-region of the pore based on mass spectrometry data from proximity labelling. This approach defined many further asymmetrically localised nuclear pore components. At the nuclear site, we identified several trypanosome-unique proteins, for instance the FG-NUPs NUP64/NUP98, but also proteins with structural homology to TREX-2 components. We mapped the components of the Ran-based nuclear export system and confirm the absence of a Dbp5 homologue. Lastly, we demonstrate, by deploying an auxin degron system, that NUP76 holds an essential role in mRNA export consistent with a functional orthology to NUP82/88.

Altogether, the combination of proximity labelling with expansion microscopy revealed an asymmetric architecture of the trypanosome nuclear pore supporting inherent roles fort directed transport. Our approach delivered novel nuclear pore associated components inclusive positional information, which can now be interrogated for functional roles to explore trypanosome specific adaptions of the nuclear basket, export control and mRNP remodelling.

## INTRODUCTION

Nuclear pores penetrate the double-membrane of the nucleus and serve as an essential gateway for the exchange of proteins, RNAs and ribosomes between the nucleoplasm and the cytoplasm. They are among the largest macromolecular complexes in nature with more than 500 copies of ∼30 different nucleoporins (NUPs) that form eight identical protomers (spokes) (Wente & Rout, 2010; Hampoelz *et al*, 2019; Schwartz, 2016; Lin & Hoelz, 2019). Each spoke is connected to the nuclear envelope (NE) as well as to the neighbouring spokes, resulting in multiple concentric rings: the inner ring (IR) at the centre of the pore is flanked by two outer rings (OR) at the cytoplasmic site (cytoplasmic ring) and nuclear site (nuclear ring). The outer rings are composed of large Y-shaped protein complexes, called the Nup84/NUP107 complex in yeast/human (Lin & Hoelz, 2019). This core structure of the pore is extended by the nuclear basket at the nuclear site and the cytoplasmic filaments at the cytoplasmic site. NUPs can be divided into three classes: (i) structured NUPs that form the scaffold of the pore, with structural features being limited to beta propellers, coiled coil and alpha-helical solenoids (ii) pore membrane proteins (POMs) that anchor the pore in the nuclear envelope via transmembrane regions and (iii) non-structured, intrinsically disordered NUPs, that contain FG (phenylalanine and glycine) repeat motifs and provide a diffusion barrier at the central channel of the pore by phase separation (Terry & Wente, 2009). While smaller molecules can pass by diffusion, the transport of larger molecules, such as most RNAs, ribonucleoprotein particles (RNPs), pre-ribosomes and most proteins, requires energy and depends on transporters. Protein transport, as well as the transport of micro-RNAs and tRNAs is mediated by importins and exportins of the karyopherin family (Wing *et al*, 2022). These transporters recognise and bind nuclear localisation signals (NLS) or nuclear export signals (NES) of their cargo and shuttle it within the phase-separated central channel of the pore by interacting with the FG-repeat NUPs (Wing *et al*, 2022). This transport is energised by the RanGTP/RanGDP gradient across the nuclear envelope maintained by the chromatin-bound guanine nucleotide exchange factor RCC1 and the cytoplasmic-localised proteins Ran-GTPase-activating protein RanGAP and RanBP1 (Wing *et al*, 2022). Importins bind cargo and Ran-GTP mutually exclusively, while exportins bind Ran-GTP and cargo cooperatively, thus allowing selective release of cargo either in the nucleus or cytoplasm, respectively, driven by GTP hydrolysis cycles of Ran (Wing *et al*, 2022). The vast majority of messenger ribonucleoprotein particles (mRNPs) are not exported by karyopherins but use the heterodimeric Mex67/Mtr2 complex (NXF1 or TAP/NXT1 in human) instead (Chen *et al*, 2024; Magistris, 2021; Ashkenazy-Titelman *et al*, 2020). The energy is provided by at least two RNA helicases, Sub2/UAP56 and Dbp5/DDX19 (yeast/human), that assemble and disassemble the Mex67/Mtr2/mRNA export complex in the nucleus and in the cytoplasm, respectively, in events known as nuclear and cytoplasmic mRNP remodelling (Xie & Ren, 2019). In ophistokonts, two asymmetric pore components, the basket and the cytoplasmic filaments, ensure directionality of RNP transport. The basket of *S. cerevisiae* consists of Nup1, Nup2, Nup60, Mlp1/2 and Pml39 and in metazoa of NUP153 (orthologue to yeast Nup1/60), NUP50 (orthologue to yeast Nup2), TPR (orthologue to Mlp1/2) and ZC3H1 (orthologue to Pml39). Nup60 anchors the yeast nuclear basket to the Y-complex of the nuclear outer ring (Stankunas & Köhler, 2024). Nup1 and NUP153 anchor the TREX-2 (three prime repair exoribonuclease 2) complex to the pore in yeast (Jani *et al*, 2014) and humans (Umlauf *et al*, 2013), respectively. TREX-2 consists of the large scaffolding protein Sac3/GANP bound to Thp1/PCID2, Sem1/DSS1, and Sus1/ENY2 (yeast/human), and, in yeast only, to Cdc31. This complex functions in recruiting the mRNP to the pore by direct interaction of the Sac3 N-terminal region with the mRNA-loaded Mex67 (Stewart, 2020). The cytoplasmic filaments are heterotrimeric complexes of Nup82/NUP88, Nup159/NUP214 and Nsp1/NUP62 (yeast/human); Nup159/NUP214 recruit the mRNA remodelling helicase Dbp5/Ddx19 with its cofactor Gle1 (Fernandez-Martinez *et al*, 2016; Bley *et al*, 2022).

Structure and composition of nuclear pore complexes has been characterised in a range of organisms (examples (Kim *et al*, 2018; Kosinski *et al*, 2016; Lin *et al*, 2016; Allegretti *et al*, 2020; Mosalaganti *et al*, 2018)). While the general structure of the pore, in particular the inner ring structure, is highly conserved, the more peripheral structures can differ between organisms and even within the same organism (Fernandez-Martinez & Rout, 2021). Yeast for example has up to three pore variants that differ in the number of nuclear outer rings and in the presence or absence of a basket (Akey *et al*, 2022; Singh *et al*, 2024; Niepel *et al*, 2005; Galy *et al*, 2004). Trypanosomes have separated from the eukaryotic main branches very early and their nuclear pore architecture is thus an important puzzle-stone towards a better understanding of pore evolution. In particular, structural differences would help to unravel which pore features constitute organism-specific adaptations and which have been present in the LECA (last common eukaryotic ancestor) (Padilla-Mejia & Field, 2023; Obado *et al*, 2017).

In *Trypanosoma brucei*, 22 NUPs were initially identified based mostly on structural similarities to human and yeast NUPs, as the sequences are poorly conserved, and pore localisation was confirmed by GFP-tagging (DeGrasse *et al*, 2009). This served as foundation for a hallmark follow-up study, that has defined the sub-complexes, quaternary structure and pore-associated proteins by a large set of immunoprecipitations with multiple baits from cryomilled samples, combined with immunogold electron localisation and *in silico* prediction tools (Obado *et al*, 2016). Similar to all other eukaryotes studied so far, the inner ring is mostly conserved (Obado *et al*, 2016; Makarov *et al*, 2021; Fernandez-Martinez & Rout, 2021), with the one exception of the membrane anchoring mechanism: *T. brucei* lacks orthologues to all POMs of opisthokonts. Instead, TbNUP65 has evolved a C-terminal transmembrane helix to connect to the nuclear envelope (Obado *et al*, 2016), replacing the amphipathic lipid-packing sensor (ALPS) motif used by its opisthokont orthologues ScNup53/59 and HsNUP35. Two additional POM candidates, with transmembrane helixes, were recently identified within the interactome of lamin-like proteins (Butterfield *et al*, 2024). The structured outer ring complex (Y-complex) was clearly defined in multiple affinity purifications to consist of TbNUP158, TbSEC13, TbNUP41, TbNUP82, TbNUP89, TbNUP132, TbNUP152 and, likely, TbNUP109 (Obado *et al*, 2016). This complex, named NUP89 complex, is the equivalent to the outer ring complexes Nup84/NUP107 from yeast/human and mostly conserved, with some lineage specific variations in the β-propeller proteins (Obado *et al*, 2016). Three FG-NUPs, NUP64, NUP75 and NUP98, are unique to trypanosomes and part of one complex with unknown localisation (Obado *et al*, 2016). There were two major unexpected outcomes from this study: (i) no asymmetrically localised NUPs were identified, with the exception of the basket proteins NUP110 and NUP92, suggested as putative homologues to yeast Mlp1 and Mlp2. Even TbNUP76, which was co-isolated with TbMex67 and has structural homology to the cytoplasmic-site specific yeast Nup82 that has a function in mRNA export (Hurwitz & Blobel, 1995; Grandi *et al*, 1995) was predicted at both outer rings by immunogold labelling; (ii) the authors could not identify any homologue to the cytoplasmic mRNA remodelling enzyme, the DEAD-box RNA helicase Dbp5. Instead, they found co-purification in high-stringency conditions between the conserved mRNA transporter TbMex67 with Ran, RanBP1 and a putative RanGAP, indicating that mRNA export may be fuelled by the Ran system. Meanwhile, many additional nuclear pore localised proteins were identified, primarily by the genome-wide localisation database TrypTag (Billington *et al*, 2023), of which most remain functionally uncharacterised.

We were puzzled by the absence of asymmetric NUPs at the outer rings, which are viewed as key determinants underpinning directed transport of macromolecules. We therefore revisited the ultrastructure of the trypanosome nuclear pore using a novel, powerful combination of expansion microscopy and proximity labelling techniques. Our approach indeed identified a set of asymmetric components and we employed these as markers to map all 75 nuclear pore localised proteins reported by TrypTag (Billington *et al*, 2023). Altogether, we provide an updated, comprehensive map of the pore and its associated proteins, including proteins of the Ran GTPase transport system. We describe many novel proteins at the nuclear site of the pore, most of these trypanosome-unique, including three potential TREX-2 complex proteins. We find the NUP76 complex proteins, NUP76, NUP140 and NUP149, exclusively at the cytoplasmic site and demonstrate a conserved function of NUP76 in mRNA export, while NUP140 and NUP149 are unique to trypanosomes, and lack any conserved binding site for Dbp5, consistent with the absence of this RNA helicase. Our data, combined with the data of (Obado *et al*, 2016), support a model of the trypanosome pore with a conserved core structure, but with a fundamental different mRNA remodelling platform at the cytoplasmic site and many trypanosome-unique proteins at the basket site that await functional characterisation.

## MATERIAL AND METHODS

### Bioinformatics

All sequences were retrieved from TriTrypDB (Shanmugasundram *et al*, 2023). InterPro was used for domain search based on sequence (Paysan-Lafosse *et al*, 2022). Homology search based in primary and predicted secondary alignments was done with Phyre2 (Kelley *et al*, 2015). Tertiary alignments of Trypanosomatid-optimised predicted AlphaFold2 models (Wheeler, 2021) were carried out with Foldseek (Kempen *et al*, 2024). Foldseek searches were performed on the web server (https://search.foldseek.com) covering all available databases (AlphaFold/Proteome v4, AlphaFold/Swiss-Prot v4, AlphaFold/Swiss-Prot v4, BMFD 20240623, CATH50 4.3.0, Mgnify-ESM30 v1, PDB100 20240101 and GMGCL 2204) with Mode 3Di/A. The outputs from Foldseek including the superimposed structures, the values of sequence identity, RMSD (root mean square deviation), TM (template modelling score), qTM and tTM (TM scores normalised by query and template length, respectively) values were retrieved. All structures were predicted using AlphaFold2-Multimer-v2.3.1 (Jumper *et al*, 2021; Evans *et al*, 2022) through the ColabFold version 1.5.3 with Mmseq2 (UniRef+Environmental) with 5 recycles and 5 models (doi:10.1038/s41592-022-01488-1). Predicted structures were visualized with ChimeraX (Meng *et al*, 2023). Heatmaps were generated with the ComplexHeatmap package (Gu, 2022) in R. The *t*-test difference values from the affinity purifications (detailed below) were fed in and clustered with a Pearson distance method (option cluster_rows = TRUE, cluster_columns = FALSE). The *t*-test difference values are represented as a colour scale and the colouring was made by the package. pLLDT plots of local prediction confidence over the protein length shown near the heatmaps are available and were retrieved from the Trypanosomatid-optimised AlphaFold2 database (Wheeler, 2021). Violin plots were created with the ggpubR package of R (https://rpkgs.datanovia.com/ggpubr/).

### Trypanosoma cells

*Trypanosoma brucei* Lister 427 procyclic cells in logarithmic growth were used for all experiments. Cells were grown in SDM-79 supplemented with 5% (v/v) FCS and 75 μg/ml hemin at 27°C, 5% CO_2_, and appropriate drugs (Brun & Schönenberger, 1979). Drugs used for transgenic cells were G418 disulfate (15 μg/ml), blasticidine S (10 μg/ml), puromycin dihydrochloride (1 μg/ml), hygromycin B (25 μg/ml) and phleomycin (2.5 µg/ml); these concentrations were used for maintenance and doubled during the actual selection process after transfection. Growth was measured by sub-culturing cells daily to 10^6^ cells/ml and measuring densities 24 hours later using a Coulter Counter Z2 particle counter (Beckman Coulter) over five days.

Transgenic trypanosomes were generated by standard procedures. Endogenous tagging with TurboID-Ty1, 3xHA, 4xTy1 and *Os*AID-3xHA was done using a PCR-based method and the pPOTv7 system (Dean *et al*, 2015). TurboID-Ty1, 3xHA and 4xTy1 customisations of the pPOTv7 were made in this work. 25 μl of PCR reaction (PrimeSTAR MAX (Takara)) was used for transfections. The PCR product was precipitated with isopropanol, washed once with 70% ethanol in a sterile hood, resuspended in 10 μl of sterile ddH_2_O and mixed with 10^7^ cells in 400 μl of transfection buffer (Burkard *et al*, 2011). Transfections were performed with Amaxa Nucleofactor IIb (Lonza Cologne AG, Germany, program X-001) using BTX electroporation cuvettes (45-0125). Cells were recovered in 25 ml SDM-79 supplemented with 20% FCS for 18 h and diluted 1:4 in 75 ml SDM-79 supplemented with 20% FCS. Relevant drugs were added and cells plated in four 24-well plates (1 ml/well). Drug-resistant populations were analysed after 10 days and confirmed with Western blotting or diagnostic PCR from genomic DNA (Rotureau *et al*, 2005). For ectopic, inducible expression of Mex67 fused to TurboID-Ty1 at the carboxy-terminus, its open reading frame was cloned in frame with TurboID-Ty1 in a genetic cassette containing a EP procyclin promoter controlled by 2x Tet operator flanked by sequences for integration to the rRNA locus (Sunter *et al*, 2012).

For Auxin-inducible degron experiments, 50 µM 5-Ph-IAA (MedChem Express, HY-134653) was added to the cultured cells from a 50 mM stock in DMSO. The auxin system was kindly provided by the laboratory of Mark Carrington (University of Cambridge, UK) and is described in (Gabiatti *et al*, 2024).

### Plasmids and PCR products

All plasmids and PCR products used in this study are listed in Table S1. pPOTv7 variants for TurboID-Ty1, 3xHA, 4xTy1 tagging were generated by sub-cloning the respective tag sequence in the BamHI/HindIII sites. pPOTv7 *Os*AID-3xHA was generated in (Gabiatti *et al*, 2024).

### Western blot and antibodies

Western blots were done using standard methods. Primary antibodies used for detection of proteins were rat anti-HA (3F10, Roche) (1:1000), anti-Ty1/BB2 ((Bastin *et al*, 1996) hybridoma supernatant 1:1000) and anti-*T.brucei* PFRA/B (L13D6) (1:10,000) (KOHL *et al*, 1999). Secondary antibodies were IRDye® 680 RD and 800 CW rat and mouse anti-goat (LI-COR) (1:30,000). Biotinylated proteins were detected with Streptavidin-IRDye® 680 LT (LI-COR) (1:10,000). Blots were scanned with the Odyssey Infrared Imaging System (LI-COR Biosciences, Lincoln, NE).

### Protein retention Expansion Microscopy (proExM) and ultrastructural Expansion Microscopy (U-ExM)

The proExM and UExM methods were performed as previously described in (Odenwald *et al*, 2024), with the following minor modifications for UExM: the primary and secondary antibody labelling reactions were done in 6-well plates with 1 ml of antibody diluted in PBS-T (0.1% Tween20 (v/v) in 3% BSA (w/v)). Primary antibodies were incubated overnight at 37°C with agitation and secondary antibodies for 3 hours at 37°C with agitation. The plates were slightly tilted to ensure that the whole gel piece was covered.

### Microscopy and quantification of microscopy data

For all fluorescence microscopy experiments, images were acquired using a fully automated iMIC microscope (TILL Photonics) equipped with a 100x, 1.4 numerical aperture objective (Olympus, Japan) and a sensicam qe CCD camera (PCO, Germany). Z-stacks (75, 100, or 150 slices, 140 nm step size) were recorded. Exposure times ranged between 50-100 ms for DAPI and 400-800 ms for all other fluorophores. Image stacks were deconvolved with the Huygens™ Essential software v24.04 (SVI, Hilversum, Netherlands). Deconvolution parameters were kept constant for all images, except for the number of iterations which were optimised depending on the signal intensity and background. To correct aberrations due to the refractive index mismatches occurring at different depths of the gel specimens, the varPSF function of Huygens Essential was used, which calculates the PSFs according to depth position. After deconvolution, images were corrected for the chromatic shift aberration between the green and red channel in three dimensions. The Huygens™ Chromatic Aberration Corrector was used with image stacks of TetraSpeck fluorescent microspheres (T7279, Thermo Fisher Scientific) as template. Fiji (Schindelin *et al*, 2012) was used for figure generation.

### Streptavidin affinity purification and LC MS/MS analysis

Affinity purification of biotinylated proteins followed by tryptic digest and peptide preparation were done as described (Moreira *et al*, 2023), except that 1 mM biotin was added to the on-beads tryptic digests, to improve the elution. Eluted peptides were analysed by liquid chromatography coupled to tandem mass spectrometry (LC-MS/MS) on an Ultimate3000 nano rapid separation LC system (Dionex) coupled to an Orbitrap Fusion mass spectrometer (Thermo Fisher Scientific).

Spectra were processed using the intensity-based label-free quantification (LFQ) in MaxQuant version 2.1.3.0 (Cox *et al*, 2014; Cox & Mann, 2008) searching the *T. brucei brucei* 927 annotated protein database (release 64) from TriTrypDB (Aslett *et al*, 2010). Analysis was done using Perseus (Tyanova *et al*, 2016) essentially as described in (Zoltner *et al*, 2020). Briefly, known contaminants, reverse and hits only identified by site were filtered out. LFQ intensities were log_2_-transformed and missing values imputed from a normal distribution of intensities around the detection limit of the mass spectrometer. A Student’s *t-* test was used to compare the LFQ intensity values between the duplicate samples of the bait with untagged control (WT parental cells) triplicate samples. The -log_10_ *p*-values were plotted versus the *t*-test difference to generate multiple volcano plots (Hawaii plots). Potential interactors were classified according to their position in the Hawaii plot, applying cut-off curves for significant class A (SigA; FDR = 0.01, s0 = 0.1) and significant class B (SigB; FDR = 0.05, s0 = 0.1). The cut-off is based on the false discovery rate (FDR) and the artificial factor s0, which controls the relative importance of the *t*-test *p*-value and difference between means (at s0 = 0 only the *p*-value matters, while at non-zero s0 the difference of means contributes). Perseus was also used for principal component analysis (PCA), the profile plots and to determine proteins with similar distribution in the plot profile using Pearson’s correlation. All proteomics data have been deposited at the ProteomeXchange Consortium via the PRIDE partner repository (Perez-Riverol *et al*, 2019) with the dataset identifiers PXD055934 (Nup75, Ran, Mex67), PXD047268 (NUP76, NUP96, NUP110) and PXD031245 (NUP158, wt control).

## RESULTS

### Expansion microscopy identifies novel asymmetric components of the trypanosome pore

We revisited the trypanosomatid nuclear pore architecture with expansion microscopy. Therefore, we expressed target proteins fused to a small peptide epitope-tag (3xHA or 4xTy1) to allow immunofluorescence detection via anti-HA or anti-Ty1. In some experiments, we expressed the target protein fused to the biotin ligase TurboID (Branon *et al*, 2018), followed by the detection of the auto-biotinylation with fluorescent streptavidin (=streptavidin imaging). We had previously shown that labelling with streptavidin increases the signal with no obvious loss in resolution, which is essential since expansion microscopy causes a massive reduction in antigen density (Odenwald *et al*, 2024). Even more importantly, streptavidin readily labels proteins within phase-separated areas, such as the nuclear pore central channel, that we found largely inaccessible to antibodies (Odenwald *et al*, 2024). Since TurboID will not only auto-biotinylate the bait but also adjacent proteins, there is the possibility that the streptavidin signal may not reflect the true localisation of the bait. Hence, throughout this work, we have confirmed all major findings derived from streptavidin labelling with orthogonal methods, such immunofluorescence, mass spectrometry and/or vice-versa labelling. All fusion proteins were expressed from the endogenous loci, to avoid changes in gene expression.

First, we used Protein Retention Expansion Microscopy (proExM), a method that expands cells after protein labelling (Tillberg *et al*, 2016). We obtained an expansion factor of 3.6 and confirmed the isotropic expansion of the nucleus (Figure S1). We initially focused on the proteins of the NUP76 complex (NUP76, NUP140, and NUP149) as these were previously shown to co-precipitate with the trypanosome mRNA export factor Mex67 (Obado *et al*, 2016). These NUPs were fused to either N- or C-terminal peptide tags (NUP76-3xHA, NUP140-4xTy1, 3xHA-NUP149). In the same cell lines, we co-expressed the nuclear basket protein NUP110 with an amino-terminal fusion to a different epitope tag (4xTy1 or 3xHA). Upon dual labelling with anti-Ty1 and anti-HA, we carried out expansion and imaging. All four proteins resolved as single dots. The signals from the NUP76-complex proteins were in all cases clearly separated from the NUP110 signal towards the cytoplasmic site of the pore (Figure 1A and Figure S2 in supplementary material). Notably, we observed for every dot signal originating from the NUP76 complex a corresponding NUP110 dot, indicating that trypanosomes, unlike yeast (Galy *et al*, 2004), do not have basket-less pores. The median, expansion-factor corrected distance to NUP110 was 129±17 nm for NUP76, 120±22 nm for NUP140, and 137±18 nm for NUP149 (Figure 1B). The large distance of the NUP76 complex proteins to the basket protein NUP110 and the absence of “double-dots” for the NUP76-complex strongly indicate asymmetric localization of the NUP76 complex exclusively at the cytoplasmic site of the pore. The data disagree with previous observations derived from immuno-electron microscopy that place NUP76, symmetrically, to both outer rings (Obado *et al*, 2016).

**Figure 1.**
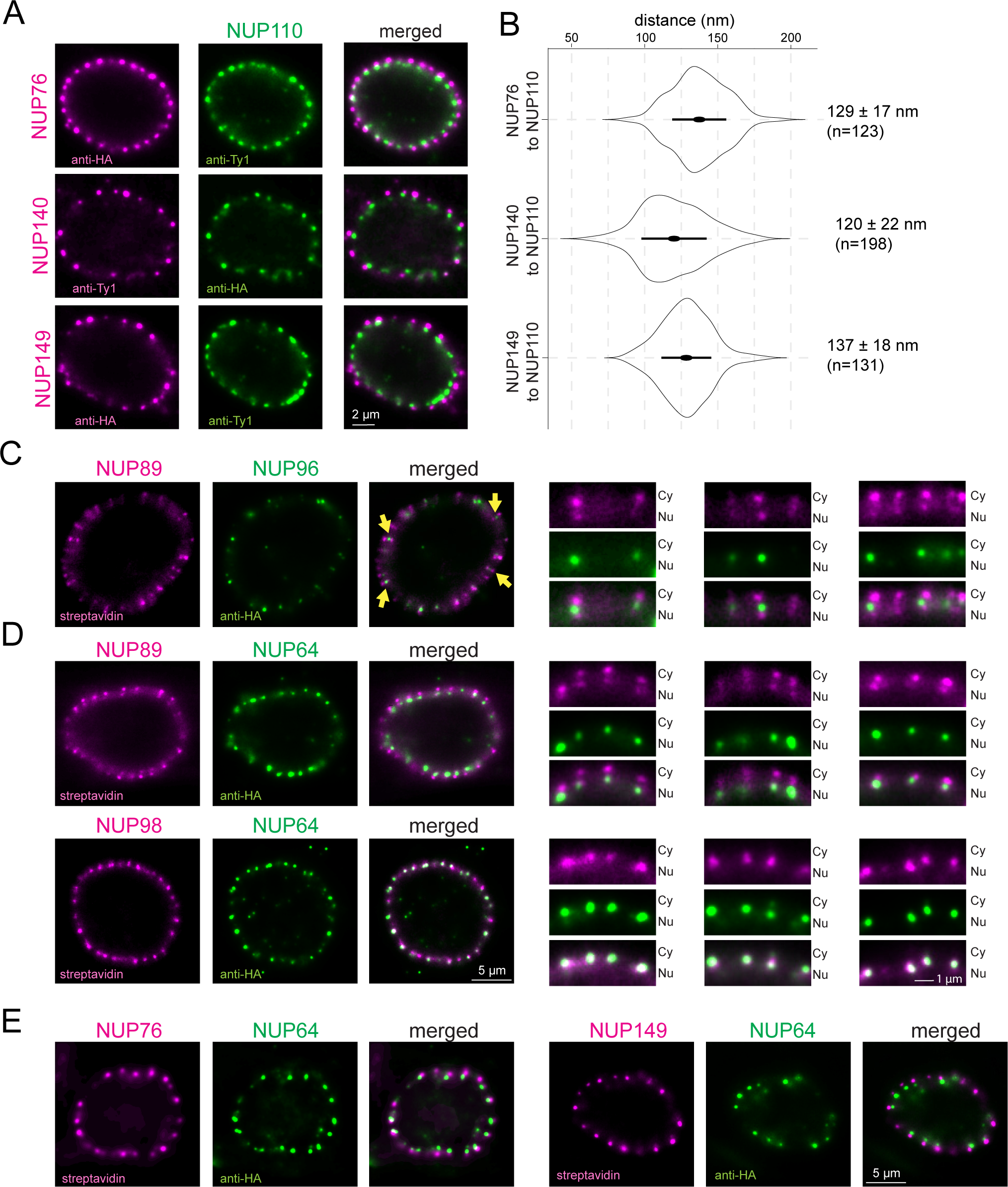
Expansion microscopy identifies novel asymmetric pore proteins. **(A)** Protein retention Expansion Microscopy (proExM) of cell lines co-expressing epitope-tagged versions of NUP76, NUP140 or NUP149 in combination with NUP110. Proteins were labelled with antibodies to the epitope tags, as indicated, before expansion. Images were deconvolved (20 iterations for NUP149 and NUP76; 60 iterations for NUP140) and single planes of the nuclei are shown. Further images are shown in Figure S2. **(B)** The distances between NUP76/NUP140/NUP149 and NUP110 were measured from manually selected pores that had both proteins in the same focal plane. The data were corrected for the expansion factor are shown as a violin plot. Mean ± standard deviation is on the right. **(C and D)** Ultrastructure Expansion Microscopy (UExM) of cells expressing proteins fused to TurboID or 3HA, as indicated. Labelling was done with fluorescent streptavidin and with anti-HA. Images were deconvolved (20, 20, 40 iterations for NUP89/NUP96, NUP89/NUP64 and NUP98/NUP64, respectively). A single plane image of an entire nucleus is shown on the left and three enlarged regions from the same or another nucleus are shown on the right (Cy=cytoplasm, Nu=nucleus). For NUP89/NUP96, yellow arrows point to pores that are in a suitable focal plane to see the two NUP89 dots sandwiching the NUP96 dot. **(E)** Ultrastructure Expansion Microscopy (UExM) of lines co-expressing TurboID-tagged versions of NUP76 or NUP149 with NUP64-3HA. Labelling was done with fluorescent streptavidin and anti-HA. Images were deconvolved with 60 and 20 iterations for NUP76/NUP64 and NUP149/NUP64, respectively. A single plane image of one nucleus is shown. Similar results were obtained for all NUP76 complex proteins (NUP76, NUP140, NUP149) with sole antibody labelling (Figure S3).

Thus, we were concerned that our observed sole cytoplasmic localisation of the NUP76 complex proteins could be a technical artifact caused by (i) an insufficient resolution of the proExM method (ii) a non-isotropic expansion across the nuclear membrane or (iii) reduced accessibilities of antibodies to the nuclear ring in comparison to the cytoplasmic ring. To investigate, we applied Ultrastructural Expansion Microscopy (U-ExM) (Gambarotto *et al*, 2019), which offers a higher resolution because the antibody labelling is applied after the expansion and the linkage error (distance between the fluorophore of the secondary antibody and the target protein) is therefore not expanded. U-ExM has been successfully used in trypanosomes (Kalichava & Ochsenreiter, 2021; Gorilak *et al*, 2021; Hernández *et al*, 2023; Odenwald *et al*, 2024), and we achieved an expansion factor of 4.2-fold with isotropic expansion of the nucleus (Figure S1). To prove that U-ExM provides the resolution to resolve the nuclear outer ring, inner ring, and cytoplasmic outer ring we first imaged NUPs that are conserved across eukaryotes. As an outer ring marker, we chose NUP89, the trypanosome orthologue to the outer ring Y-complex component of yeast Nup84/85 and human NUP107/75 (Obado *et al*, 2016). As an inner ring marker, we selected NUP96, the conserved trypanosome orthologue to *S. cerevisiae* Nic96. We co-expressed NUP89-TurboID with NUP96-3HA. U-ExM with fluorescent streptavidin and anti-HA readily resolved NUP89 as a double dot, sandwiching the single dot signal of NUP96 (Figure 1C), proving that the resolution of the method is sufficient to distinguish these different subregions of the pore. However, as the NUP96 signal was weak, we searched for a better inner ring marker. We tested NUP64, a trypanosome-unique FG-repeat NUP previously identified as a multi-complex NUP localised mostly to the centre of the pore (Obado *et al*, 2016). To our surprise, the resulting single NUP64 dot signal was not sandwiched by the two outer ring dots of NUP89 but instead co-localised solely with the NUP89 dot at the nuclear site (Figure 1D). Likewise, TbNUP98, known to form a complex with NUP64 (Obado *et al*, 2016), resolved as a single dot that colocalised exclusively with the NUP64 dot at the nuclear site (Figure 1D). The data indicate an asymmetric, exclusive nuclear-site localisation of NUP64 and NUP98 (Figure 1D).

Having confirmed that UExpM readily resolves the two outer rings, we reassessed the localisation of the NUP76 complex. Using identical cell lines as for ProExM (Figure 1A) we obtained the same result: all NUP76 complex proteins were resolved as single dots at the cytoplasmic site, clearly separated from NUP110 (Figure S2). We further confirmed the sole cytoplasmic localisation of the NUP76 complex by co-staining C-terminal fusions of this complex to either 3xHA (Figure S3) or TurboID (Figure 1E) with a C-terminal 4xTy1 fusion of our newly-identified nuclear-site marker NUP64.

In summary, we discovered five asymmetric proteins of the trypanosome nuclear pore complex, previously assumed to be symmetrically distributed: NUP76/NUP140/NUP149 at the cytoplasmic site and NUP64/NUP98 are at the nuclear site. With the exception of NUP76, which is the structural orthologue to yeast NUP82 and human NUP88 (Obado *et al*, 2016), all novel asymmetric NUPs are trypanosome-specific.

### A proximity map of the trypanosome nuclear pore

The novel availability of asymmetric NUPs prompted us to use mass spectrometry data from TurboID proximity labelling experiments, to generate a proximity map of the entire trypanosome nuclear pore. We used six available LC-MS/MS datasets from streptavidin-affinity purifications, namely amino- and carboxy-terminal TurboID fusions of NUP110 (basket), NUP96 (inner ring) and NUP76 (cytoplasmic outer ring) (Odenwald *et al*, 2024). Of all the proteins that were labelled by these baits, we initially focused on bona fide NUPs. For each dataset, we colour-coded the enrichment of the NUP proteins based on *t*-test difference to a wild-type control. Then, the NUPs were sorted by hierarchical clustering applying a Pearson distance method (Figure 2A). We included the pLLDT plots (Wheeler, 2021) to indicate predicted structured and unstructured (mostly FG-repeat) domains for each of the NUPs. The majority of the NUPs separated into three clearly distinct main clusters. The first cluster contained the basket proteins (NUP110, NUP92), the lamina protein NUP2 (Maishman *et al*, 2016) and NUP64/NUP98 that we had placed asymmetrically localised to the nuclear site by expansion microscopy (Figure 1D). The second cluster largely consisted of NUPs previously identified as inner ring NUPs (Obado *et al*, 2016) while the third was dominated by NUPs previously assigned to the outer ring (Obado *et al*, 2016). Eight NUPs were not or poorly labelled by the three bait proteins, either, due to their small size (Moreira *et al*, 2023) or for unknown reasons. These were manually added to the tree, using positional information from (Obado *et al*, 2016), except for NUP48 and Gle2 which could not be unequivocally assigned. For the vast majority of NUPs, the proximity map confirmed the previous assignments of the NUPs from affinity capture experiments (Obado *et al*, 2016).

**Figure 2.**
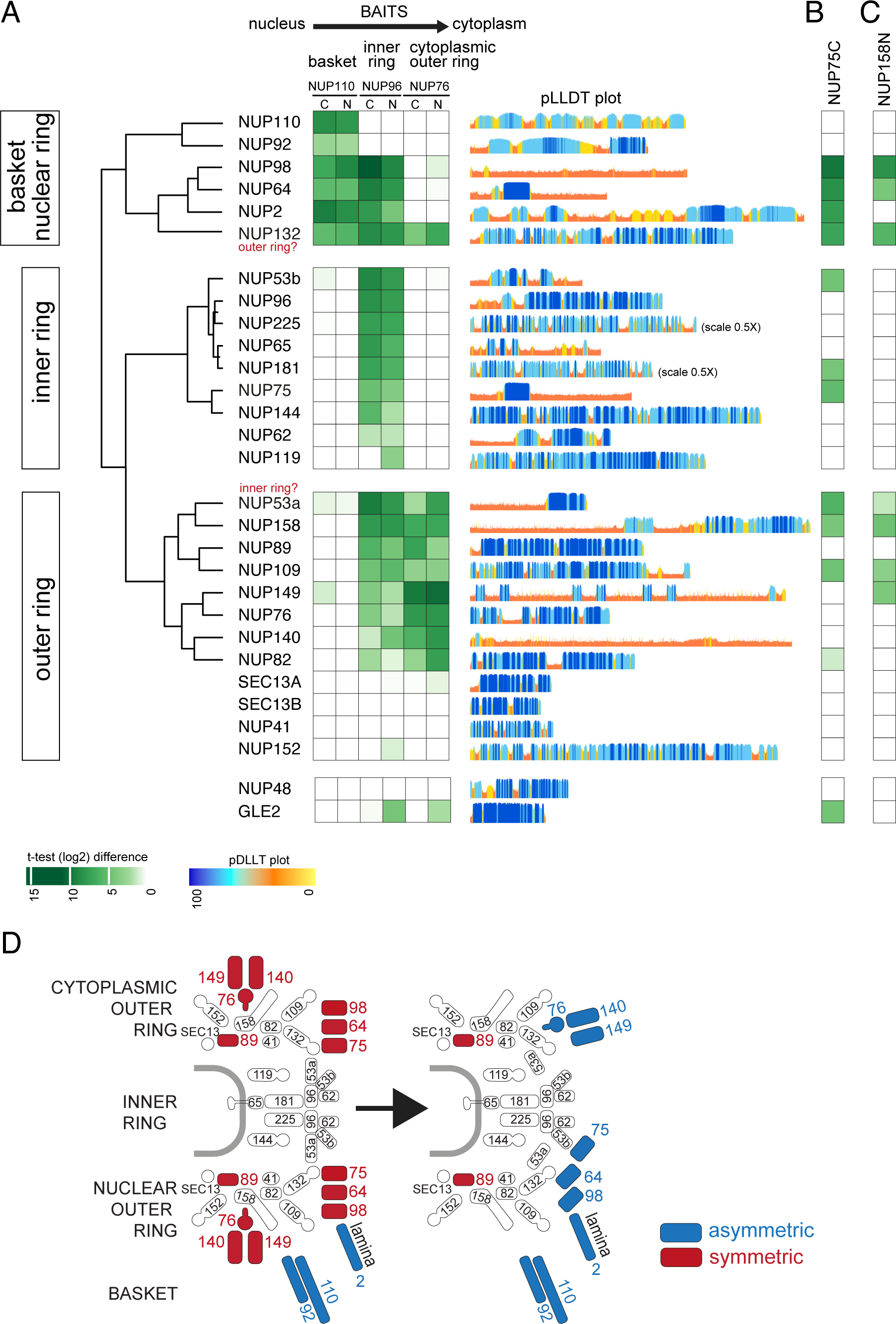
A proximity map of the trypanosomatid nuclear pore. **(A)** Mass spectrometry data (t-test difference values) from proximity labelling experiments were used to create a heat map of the trypanosome nuclear pore. N- and C-terminally tagged TurboID fusions of NUP110 (basket), NUP96 (inner ring) and NUP76 (cytoplasmic outer ring) served as baits and the labelling (proximity) of all bona fide nuclear pore proteins was analysed. The heat map grouped most NUPs into three main clusters, corresponding to the basket/nuclear ring, inner ring and outer ring. Some NUPs with too little or no labelling were manually added using data of (Obado *et al*, 2016). Only two NUPs appear potentially misplaced (NUP132 and NUP53a) and two NUPs could not be placed (NUP48 and GlLE2). pLDDT plots from Trypanosomatid-optimized AlphaFold2 models (Wheeler, 2021) are shown on the right. **(B and C)** Mass spectrometry data (t-test difference values) from proximity labelling experiments of NUP75-TurboID and TurboID-NUP158 are shown for all the NUPs shown in A, using the same colour-code. **(D)** The model of the trypanosome nuclear pore changes, with the discovery of five novel asymmetrically localised proteins.

NUP98 and NUP64 unequivocally grouped with the basket/nuclear site (basket and inner ring) and were labelled by NUP110 and NUP96 baits, but not with the cytoplasmic NUP76, in line with our proExM and U-ExM data (compare Figure 1D). To our surprise, NUP75, which shares 46% sequence identity with NUP64 and associates with both NUP98 and NUP64 (Obado *et al*, 2016), was placed to the inner ring and was not labelled by NUP110. We did a *vice versa* TurboID experiment with NUP75 as bait (Figure S4, Table S2) and confirmed the absence of NUP110 labelling, as well as the proximity of NUP75 to NUP98 and NUP64 (Figure 2B). Moreover, our previously published TurboID data with the outer ring NUP158 (carboxy-terminal fusion) strongly labelled NUP64 and slightly less NUP98, but not NUP75 (Figure 2C), further supporting the absence of NUP75 from the outer rings (Odenwald *et al*, 2024). Our data strongly suggest a model of the asymmetric NUP98/64/75 complex reaching from the nuclear outer ring to the inner ring, with NUP98 and NUP64 being located at the outer nuclear ring and NUP75 at the inner ring. Data from previous affinity isolation experiments with NUP98, NUP64 and NUP75 show marked differences between the interactomes of NUP98/64 and NUP75, including the exclusive absence of NUP110 from the NUP75 interactome, in full agreement with our data (Obado *et al*, 2016).

Only two NUPs exhibited ambiguous placement in the proximity map. The outer ring NUP132 is labelled strongly by all eight baits arguing against a sole localisation to the basket/nuclear site and suggesting a symmetric distribution with the two protomers extending to both sites of the pore. Further, the presumed inner ring NUP53a is labelled by the outer ring NUP76. However, labelling by NUP96 is stronger and the outer ring NUP158 bait labels NUP53a very weakly, indicating that NUP53a is mostly at the inner ring, potentially reaching out to the outer ring.

In summary, the proximity map accurately predicts the localisations for the vast majority of NUPs. Importantly, it offers orthogonal validation of the asymmetric localisation of NUP98 and NUP64 to the nuclear site of the pore as derived from expansion microscopy. Therefore, the NUP110/96/76 proximity map has the potential to predict localisation of proteins with sub-pore size resolution, which prompted us to look at all 44 proteins that have nuclear pore localisation according to TrypTag (Billington *et al*, 2023) but are not annotated as *bona fide* NUPs.

### Mapping the Ran-based mRNA export system to the pore

First, we focused on all nuclear-pore localised proteins that are involved in mRNA export: Mex67 (Schwede *et al*, 2009) , Mtr2, Mex67b (Obado *et al*, 2022b) and, as postulated (Obado *et al*, 2016, 2022a), Ran, RanGAP, the putative RanGDP importer NTF2 and two Ran binding proteins, RanBP1 and RanBPL (Brasseur *et al*, 2014). The proximity map places RanGAP, RanBP1 and Ran to the cytoplasmic site of the pore, while RanBPL is predicted at the nucleoplasmic site (Figure 3A). For the mRNA transporters Mex67 and, in particular for Mex67b, the labelling was weaker and less confined to a specific site. For Mtr2 and NTF2 we obtained no labelling, likely due to their small size which is problematic in BioID (Moreira *et al*, 2023).

**Figure 3.**
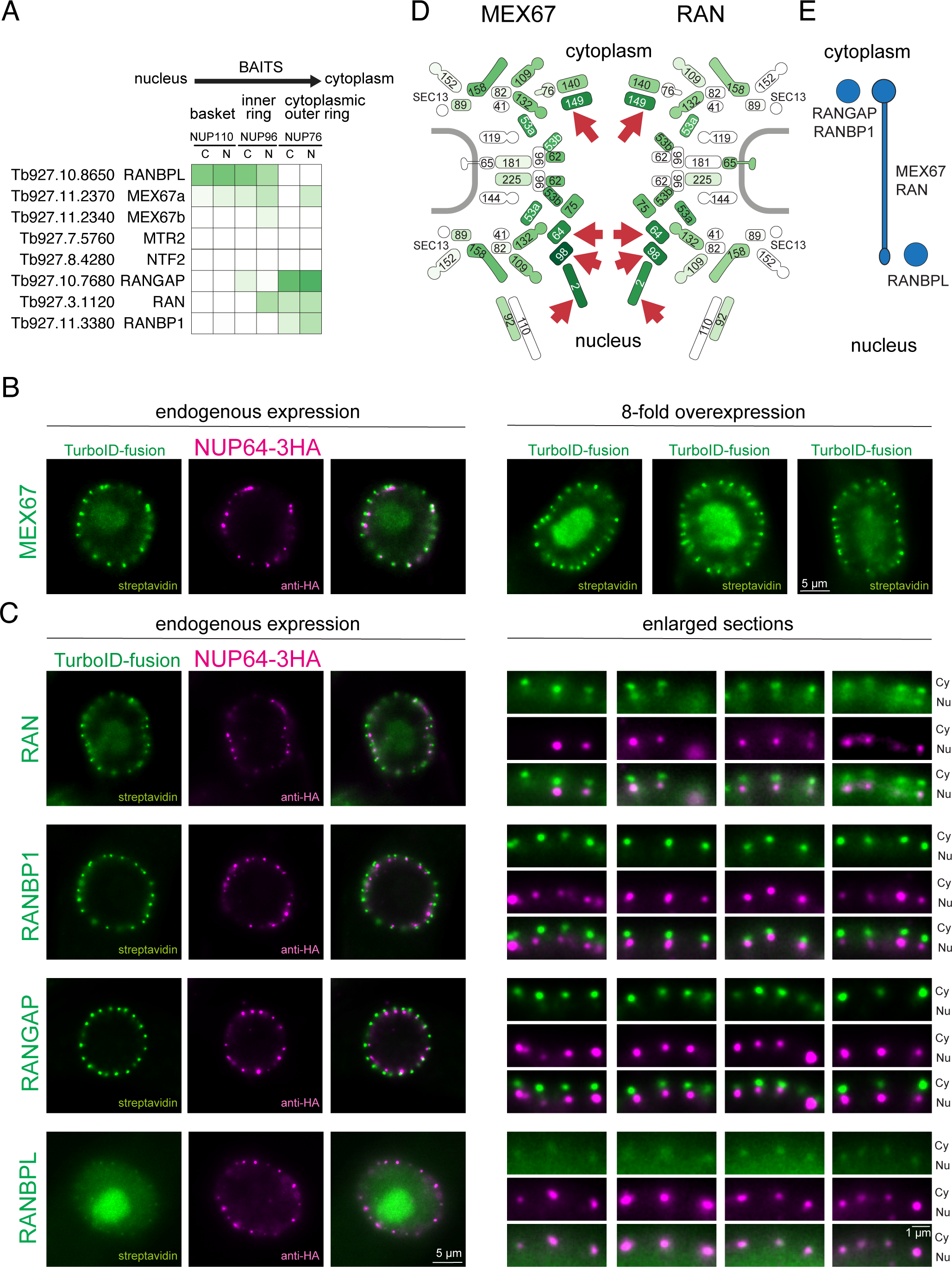
Mapping the trypanosome mRNA export system to the nuclear pore. **(A)** Mass spectrometry data (t-test difference values) from proximity labelling experiments with NUP110/NUP96/NUP76 were used to map the Ran-based mRNA export system to the trypanosome pore. **(B and C)** Ultra-expansion microscopy. Mex67, Ran, RanBP1, RanGAP and RanBPL were expressed as C-terminal (Mex67, Ran RanBP1, RanGAP) or N-terminal (RanBPL) fusions to TurboID in a cell line co-expressing the nuclear-site marker NUP64-3HA; all from the endogenous locus. Cells were expanded and the proteins detected with streptavidin and anti-HA. Images of single nuclei are shown (single plane of deconvolved Z-stacks with 10 iterations for Mex67, Ran, RanBP1 and RanBPL and 20 iterations for RanGAP). For Mex67, the streptavidin signal was weak and three representative nuclei of an overexpression cell line are shown (B, right). For the other proteins, enlarged section of the nuclear envelope of the same or other nuclei are shown (C, right). **(D)** Proximity labelling of the nuclear pore by Mex67 and Ran. Mex67 and Ran were expressed as carboxy-terminal fusions to TurboID from the endogenous loci and biotinylated peptides were analysed by mass spectrometry. The labelling of all bona-fide NUPs by Mex67 (left) and Ran (right) is shown coloured based on their *t*-test difference values in comparison to wild type cells. Asymmetric NUPs are marked with a red arrow. **(E)** Schematic summary of the UExM localisation data of the Ran-based mRNA export system.

Next, we determined the localisation of Mex67, Ran, RanGAP, RanBP1 and RanBPL by U-ExM, expressing TurboID fusions in a cell line that expressed NUP64-3xHA as a nucleoplasmic-site marker (Figure 3B, C). RanGAP and RanBP1 resolved as a single dot, well distanced from the NUP64 dot towards the cytoplasmic site, while the RanBPL signal overlapped with the NUP64 signal at the nuclear site, in full agreement with the proximity map. The biotinylation signal of the two proteins with suspected shuttling activity, Ran and Mex67, resolved as a large cytoplasmic dot and a smaller nuclear dot, connected by a string-like signal reaching through the pore. For Mex67, we observed that these bone-shaped signals were more pronounced when the gene was 8-fold overexpressed from an ectopic locus (Figure 3B, images on the right) which only slightly impaired growth (Figure S5). For Ran, Mex67 and RanBPL, we observed an additional signal at the nucleolus, which is defined by the reduction in DAPI stain (Figure S6A), and a minor signal in the nucleoplasm. For RanBPL, the nucleolar signal was stronger than the signal at the pores, while for Ran and Mex67 (at endogenous expression levels) the nuclear pore signal was dominant. We attempted to confirm the nucleolar localisation by direct immunofluorescence instead of streptavidin imaging. The nucleolus is challenging to label with antibodies due to phase separation (Odenwald *et al*, 2024), but for Mex67 fused to 4Ty1 we could get a weak, but distinct nucleolar antibody signal (Figure S6B). The functional implications of the nucleolar localisation of Ran, Mex67 and RanBPL is not fully understood in trypanosomes but not unexpected, as in ophistokonts Ran and Mex67 participate in pre-ribosome transport (Li *et al*, 2023).

For the shuttling proteins Ran and Mex67 we confirmed the streptavidin-based imaging data by LC-MS/MS upon streptavidin enrichment (Figure 3D, Figure S4, Table S2). Both Mex67 and Ran strongly labelled FG-NUPs lining the inner pore channel, consistently reflecting their movement across the pore. The NUPs with the strongest labelling were the asymmetric NUPs on both sides of the pore: NUP149 at the cytoplasmic site and NUP98, NUP64 and the lamin-like protein NUP2 at the nuclear site (red arrows in Figure 3D). This strongly suggests that both proteins, Mex67 and Ran, would have binding sites at both sides of the trypanosome pore, analogous to human Ran (Wu *et al*, 1995; Mahajan *et al*, 1997; Lindsay *et al*, 2002; Nakielny *et al*, 1999). There was weak labelling of structured NUPs, explaining why the NUP76/96/110 proximity map did not resolve their shuttling (Figure 3A). Consistently, preferential labelling of asymmetric FG NUPs over structured NUPs has also been shown for human karyopherins tagged with the biotin ligase BirA* (Mackmull *et al*, 2017).

In conclusion, the NUP110/96/76 proximity map predicted the localisation of all non-shuttling Ran system components confidently and in agreement with streptavidin imaging in U-ExM. RanGAP is at the (expected) cytoplasmic site, together with RanBP1. RanBPL had not been previously mapped, but is unequivocally placed to the nuclear site, consistent with its binding preference for RanGTP (Brasseur *et al*, 2014). The NUP76/96/110 proximity map failed to categorise the shuttling proteins Mex67 and Ran, likely because key interactions with non-structured FG-NUPs are not represented in our bait repertoire. Direct BioID combined with orthogonal assessment through expansion microscopy, was required to confidently place these putative export factors. The derived localisations are summarised in Figure 3E and are consistent with a mechanistically divergent Ran-dependent mRNA export in trypanosomes.

### Mapping unknown proteins to the pore

To predict the localisation of the remaining 38 nuclear pore-localised proteins more accurately, we extended the NUP76/96/110 map by the proximity labelling data of Mex67 and Ran and of the outer ring protein NUP158 (Moreira *et al*, 2023).

Fifteen of the 38 nuclear pore-localised proteins are karyopherins (Figure S7), five of which have not been previously classified as karyopherins but have unequivocal structural homology to importin and exportin-like folds predicted by FoldSeek (Figure S7B); these include a putative orthologue to the importin Hikeshi (Tb927.1.1400) that is specialised on the import of Hsp70-family proteins (Kose *et al*, 2012) (Figure S7C). Karyopherins were mostly not, or poorly labelled by NUP76/96/110 (Figure S7A). The likely reason is their preferred interaction with FG-NUPs rather than structured NUPs, similar to what we observed for Mex67 and Ran (compare Figure 3A). The one exception is Xpo1 (exportin 1), known to be involved in the transport of both mRNAs and tRNAs (Buhlmann *et al*, 2015), which is labelled by all six bait proteins.

An additional five proteins with nuclear pore-localisation were not labelled by either of the six proteins (Figure S8). For three of these proteins, Tb927.11.1000, Tb927.10.12200 and Tb927.10.8160, the reason could be failed detection due to small size (Moreira *et al*, 2023). None of these small proteins has homologues outside of trypanosomatids and their function is unknown. Tb927.10.8160 has the strongest nuclear pore localisation ((Billington *et al*, 2023), Figure S8B) and high-throughput phenotyping indicates an essential function (Alsford *et al*, 2011). The two larger proteins (Tb927.1.3230 and Tb927.9.12700) do not have very prominent nuclear pore localisation ((Billington *et al*, 2023), Figure S8B). For Tb927.9.12700, biochemical data indicate glycosomal localisation (Güther *et al*, 2014) and Tb927.1.3230 could be the trypanosomatid ortholog of the ribosome biogenesis factor Rix7 (Lo *et al*, 2019). Their lack of labelling might thus be due to poor or absent nuclear pore localisation.

The remaining 18 proteins were labelled by at least one of the six bait proteins (Figure 4A). Strikingly, none were labelled by the cytoplasmic-site marker protein NUP76, suggesting absence of further proteins with exclusive cytoplasmic localisation, other than the NUP76 complex, RanGAP and RanBP1. Moreover, not a single protein was exclusively labelled by the outer ring NUP158 (Moreira *et al*, 2023), with the one exception of Tb927.11.13080. The (near) absence of combined labelling by NUP76 and NUP158 suggests that the outer-ring proteome may be complete. Instead, these 18 proteins were either labelled exclusively by NUP110 (7 proteins) or NUP96 (4 proteins) or by both, NUP110 and NUP96 (3 proteins). Four proteins were not labelled by NUP110/96/76, but showed some labelling with Mex67, Ran or NUP158.

**Figure 4.**
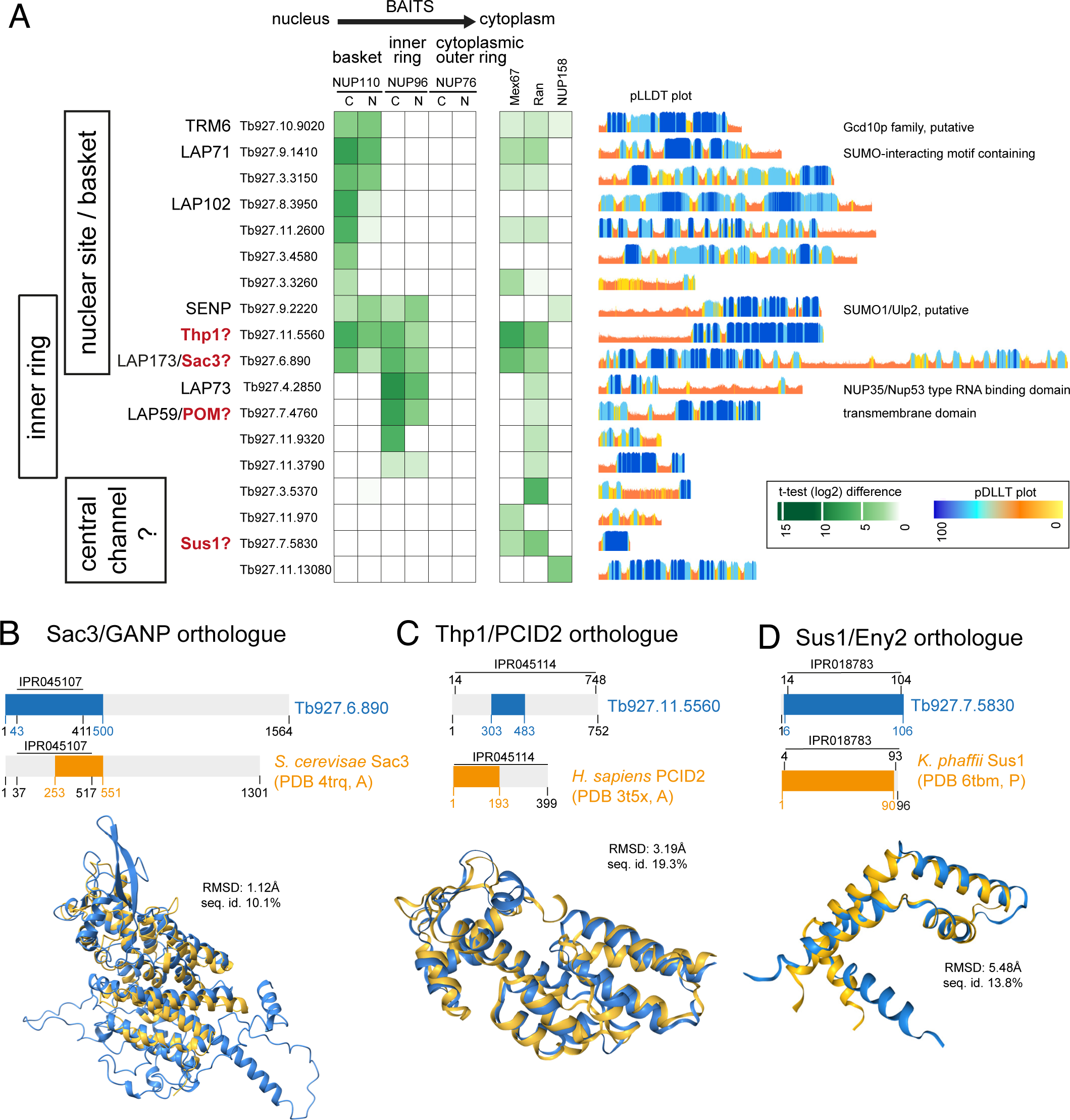
Characterisation of unknown nuclear pore proteins and their localisation over the nuclear pore complex. **(A)** Mass spectrometry data (t-test difference values) from proximity labelling experiments with NUP110/NUP96/NUP76/Mex67/Ran/NUP158 were screened for labelling of nuclear pore localised proteins, that are not bona fide NUPs or karyopherins. All proteins that were labelled by at least one of the baits proteins are shown. Annotations are explained in the text. pLDDT plots from Trypanosomatid-optimized AlphaFold2 models (Wheeler, 2021) are shown on the right. **(B-D)** Structural alignments of AlphaFold2 models of the *T. brucei* TREX2 complex candidates Sac3 (B), Thp1 (C) and Sus1 (D) with PDB structures of the respective TREX2 complex proteins from other organisms, using Foldseek (Kempen *et al*, 2024). The regions of the proteins that were used for the structural alignments are shown in the schematics in orange (*T. brucei*) and blue (other organisms). The root mean square deviation of atomic positions (RMSD) and the sequence identity are shown for the superimposed regions.

The majority of these 18 proteins is unique to Kinetoplastida or even to Trypanosomatida and lack functional annotation. Only two proteins have readily identifiable homologues outside Kinetoplastida: Tb927.10.9020 has homology to the non-catalytic, tRNA substrate binding subunit of the tRNA methyltransferase Trm6/Gcd10, responsible for adenosine(58)-N(1) methylation, a modification present in many eukaryotic tRNAs (Anderson *et al*, 2000). The *T. brucei* protein exhibits strong nuclear pore localisation (Billington *et al*, 2023) and clusters with the basket in our proximity map, which is in contrast to the nuclear localisation observed for *S. cerevisiae* and *A. thaliana* Gcd10/TmR6 (Anderson *et al*, 1998; Tang *et al*, 2020). Whilst localisation to the nuclear pore and/or envelope is not unheard of for tRNA modifying enzymes (Simos *et al*, 1996; Clark & Abelson, 1987), this finding requires further investigation, as, conversely, the putative *T. brucei* homolog (Tb927.11.11660) of the corresponding catalytic subunit, TRM61/Gdc14, localises to the nuclear lumen/nucleoplasm localisation (Billington *et al*, 2023). The second protein with homologies outside Kinetoplastida, Tb927.9.2220, is a SUMO protease of the Ulp/SENP (ubiquitin-like protease/sentrin-specific protease) family with potential function in resolving stalled DNA replication forks (Kramarz *et al*, 2020). Both the SENP protease Ulp1 of yeast and SENP2 of human have nuclear pore localisation (Hang & Dasso, 2002; Nie & Boddy, 2015) and the latter was localised to the nuclear site of the pore, consistent with our map (Hang & Dasso, 2002).

The remaining 16 proteins contain five proteins with predicted basket or inner ring localisation that were previously identified as lamina associated proteins (LAPs), based on their interactions with the lamina-like proteins NUP1 and NUP2 (Butterfield *et al*, 2024). Two of these LAPs, LAP71 and LAP102, are basket specific in our map, as expected for lamina associated proteins. Two further LAPs, LAP73 and LAP59, have exclusive inner ring prediction. Consistently, LAP59, conserved across eukaryotes, has two N-terminal transmembrane domains and was suggested to be a POM (pore membrane protein) (Butterfield *et al*, 2024). LAP73 has a divergent NUP35/Nup53 type RNA binding domain (Butterfield *et al*, 2024) and the *T. brucei* Nup53 ortholog, TbNup65, is anchored to the nuclear envelope via a trans-membrane helix (Obado *et al*, 2016). This raises the possibility of a nuclear envelope- and thus inner ring localisation of LAP73 via binding to TbNUP65. However, immunoprecipitation assays failed to establish an inner ring association with LAP73 (Obado *et al*, 2016). It is possible that the interaction is weak and thus exclusively detectable in BioID, as our data strongly suggests inner ring proximity, consistent with LAP73 potentially bridging NUP65 and the inner ring, a hypothesis that deserves further investigation.

Of significant interest is basket/inner ring-predicted LAP173, which has a Sac3/GANP domain and was suggested to be the orthologue to Sac3 and sole representative of a potential trypanosome TREX-2 complex (Butterfield *et al*, 2024). Association with Mex67 was observed with two methods: affinity purifications (Obado *et al*, 2016) and BioID (Moreira *et al*, 2023). In fact, the Sac3/GANP region of a LAP173 model predicted by a trypanosome-optimised AlphaFold2 (Jumper *et al*, 2021; Wheeler, 2021) displays remarkable structural homology to the equivalent region of an experimentally resolved *S. cerevisiae* Sac3 structure (RMSD 1.12Å, (Schneider *et al*, 2015)), despite poor sequence conservation (Figure 4B).

Motivated by the presence of putative Sac3, we used Foldseek (Kempen *et al*, 2024) to search for structural homologues of the remaining TREX2-complex components, using AlphaFold2 models as inputs (Jumper *et al*, 2021; Wheeler, 2021). We identified the Csn12-like domain containing protein Tb927.11.5560 as a putative Thp1 orthologue (Figure 4C), with structural homology to the human Tph1 homologue PCID2 (PDB entry 3T5X; TM score 0.82), while the primary sequence is, again, poorly conserved. Just like Sac3, our proximity map places this Thp1 candidate to both, basket and inner ring. Moreover, we identified Tb927.7.5830 as a putative Sus1 orthologue with highest structural similarity to Sus1 of the yeast *K. phaffii* (Papai *et al*, 2020) (Figure 4D). The Sus1 candidate protein is not labelled by the structural NUPs, presumably due to its small size. However, all three trypanosome TREX2 complex candidates, including Sus1, are strongly labelled by Mex67, a prototypic Sac3 interactor in ophistokonts (Ellisdon *et al*, 2012) and Ran, supportive of a potential role in a trypanosome TREX2 complex (Figure 4A). Two further Trex-2 components, Sem1 and Cdc31 (Stewart, 2020), were not identified by our modelling approach, that encompassed all nuclear pore localised proteins. Sem1 appears to act as Thp1/Sac3 complex stabilising element and also associates with other, functionally unrelated, complexes (Ellisdon *et al*, 2012). Cdc31, like Sus, binds to the Sac3 CID domain and both these factors are reported to promote nuclear basket association (Jani *et al*, 2009). It is conceivable that Cdc31 and Sem1 may be dispensable in a divergent trypanosome TREX2 complex, which warrants functional investigation.

In conclusion, our extended NUP110/NUP96/NUP76/Me67/Ran proximity map granted mapping the majority of nuclear pore localised proteins to a sub-region of the pore. We found no evidence for the existence of any further proteins asymmetrically distributed to the cytoplasmic-site other than the NUP76 complex, RanGAP and RanBP1, indicating completeness of the cytoplasmic-site specific proteome of the pore. Instead, we predict a diverse cohort of proteins with preferential localisation to the basket or nuclear site of the pore, including three putative TREX2 complex proteins with proximity to Ran and Mex67, indicative of a conserved function. These candidate TREX2 components, as well as the other newly assigned basket-specific proteins now await experimental analysis, to understand the mechanistic details of the trypanosome mRNA export platform.

### The trypanosome NUP76 complex as a cytoplasmic mRNA remodelling hub

We have assigned the NUP76 complex (NUP76, NUP140, NUP149) exclusively to the cytoplasmic site (Figure 1A, E) and a previous study has shown the interaction of this complex with Mex67 under high stringency conditions (Obado *et al*, 2016). In combination, these data strongly suggest that the NUP76 complex is the trypanosomes cytoplasmic mRNP binding hub that serves as remodelling platform. In opisthokonts, the cytoplasmic mRNP remodelling platform is based on the heterotrimeric complex composed of Nup82/NUP88, Nup159/NUP214 and Nsp1/NUP62 (yeast/human) (Figure 5A) (Fernandez-Martinez *et al*, 2016; Bley *et al*, 2022). The three proteins are connected via a carboxy-terminal parallel coiled-coil structure. Both Nup82/NUP88 and Nup159/NUP214 possess N-terminal β-propellers, that provide direct binding platforms for Nup145N/NUP98 (which recruits Gle2/RAE1) and the RNA helicase Dbp5/DDX19 (recruiting Gle1/GLE1), respectively, (yeast/human, Figure 5A). Nsp1/NUP62 and Nup159/NUP214 possess FG repeat regions. *T. brucei* NUP76 has been previously suggested as Nup82/NUP88 homologue (Obado *et al*, 2016); indeed, the AlphaFold2 model of NUP76 (Jumper *et al*, 2021; Wheeler, 2021) shows an analogous structural organisation with a β-propeller at the amino-terminus, interrupted by a long, disordered coil, and a three-helical coiled-coil at the carboxy-terminus (Figure 5B-C and Figure S9). Moreover, *T.brucei* NUP76 may share its β-propeller interactions with NUP82/88: orthologues to both Nup145N/NUP98 and Gle2/RAE1 can be readily identified in trypanosomes (DeGrasse *et al*, 2009; Obado *et al*, 2016). However, the two remaining TbNUP76 complex components, TbNUP140 and TbNUP149, do not exhibit detectable structural homology to the NUP82/NUP88 partner proteins Nup159/NUP214 and Nsp1/NUP62 (Figure 5A-C) as based on AlphaFold2 predictions. TbNUP140 consists almost entirely of FG repeats of the PxFG type, apart from a small N-terminal stretch with a coiled-coil structure. predicted with low confidence. NUP149 is not FG rich, with only few FG motifs of the SxFG and of the FxFG type but contains six zinc fingers sparsed by coils and potentially a small coiled-coil region at the carboxy-terminus (Obado *et al*, 2016). Notably, zinc fingers are also present in the human cytoplasmic-filament NUP358 and the nuclear-site localised NUP153, both absent from trypanosomes, where they engage in Ran binding (Yaseen & Blobel, 1999; Higa *et al*, 2007). However, the zinc fingers of TbNUP149, confidently predicted as three β-hairpin strands with four cysteines side chains coordinating a zinc ion by AlphaFold2 and AlphaFold3 (Abramson *et al*, 2024) appear to lack obvious sequence or structural homology to the zinc fingers of human NUP358 and NUP153 (Figure S10). Whether these TbNUP149 zinc fingers promote Ran binding, analogous to human NUP358 and NUP153, remains to be investigated. Notably, NUP149 is heavily biotinylated by Ran-TurboID, indicative of a possible interaction (Figure 3B).

**Figure 5:**
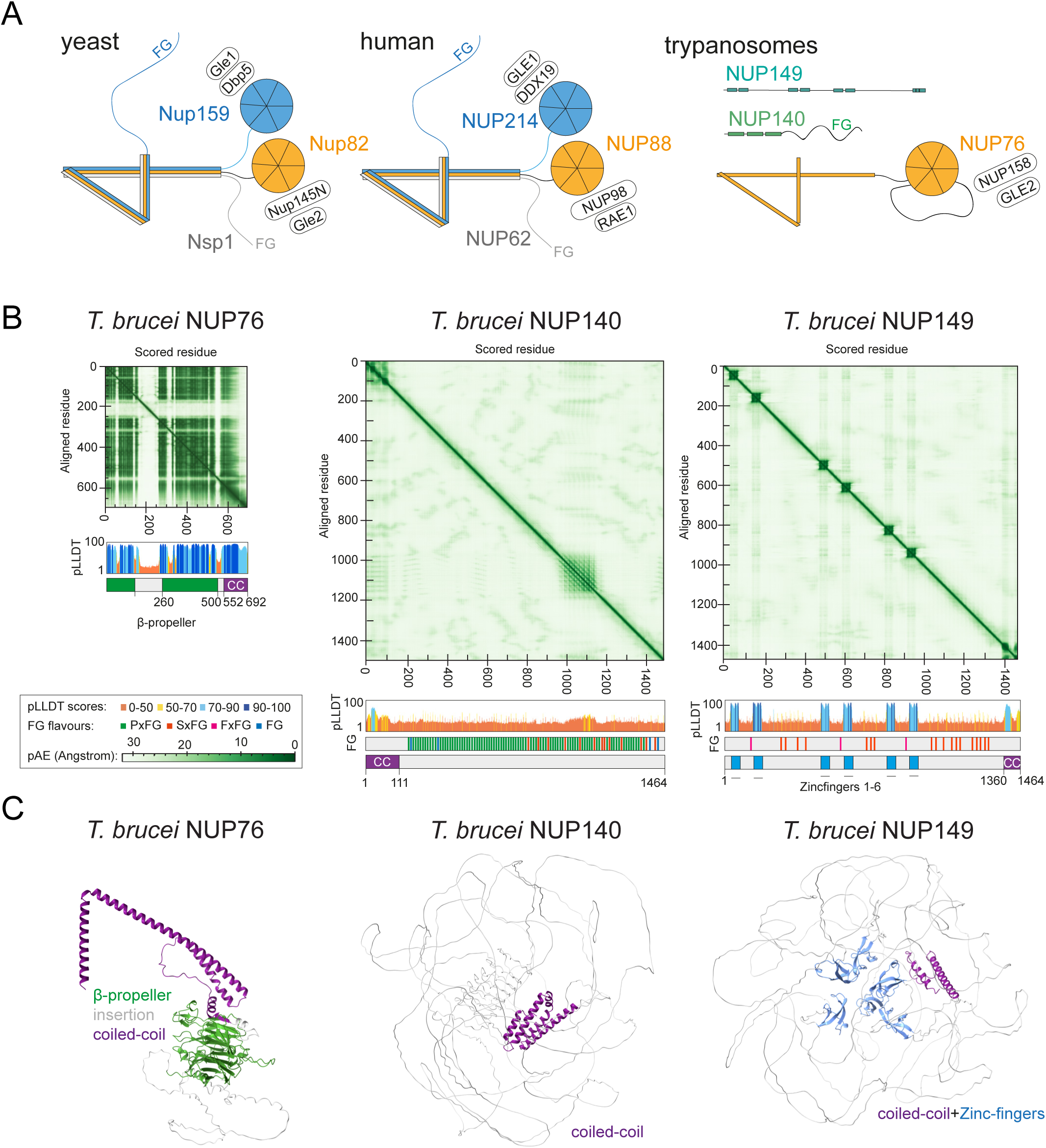
The *T. brucei* NUP76 complex is only partially conserved. **(A)** Schematics of the cytoplasmic filament complex from yeast and human (modified from (Bley *et al*, 2022), not to scale. The proteins of the trypanosome NUP76 complex are shown for comparison (left). Note that trypanosomes do have orthologues to NUP145N and Gle2, but it is not known whether these directly interact with NUP76. **(B)** pAE and pLLDT plots of trypanosomatid-optimised AlphaFold2 models of NUP76, NUP140 and NUP149. Each protein is also shown schematically with all predicted domains and, for NUP140 and NUP149, with positions and types of FG repeats. **(C)** Models of trypanosomatid-optimised AlphaFold2 predictions of NUP76, NUP140 and NUP149. Structured parts are coloured, disordered regions are shown in grey.

To investigate whether the trypanosome NUP76 complex is involved in mRNA export, we depleted the protein with an auxin-inducible degron system. Both alleles of the NUP76 gene were genetically fused to the OsAID-3HA sequence, in a cell line that expressed the necessary components for the auxin degron system (Gabiatti *et al*, 2024); the cell line was confirmed by diagnostic PCR (Figure S11). Upon induction with the auxin derivative 5-Ph-IAA, the NUP76-OsAID-3HA protein was depleted within 2 hours (Figure 6A), followed by growth arrest (Figure 6B) and accumulation of poly(A) signal in the nucleus that was saturated 4 hours post induction (Figure 6C). This phenotype is similar to the one observed upon Nup82 depletion in yeast (Hurwitz & Blobel, 1995; Grandi *et al*, 1995), strongly suggesting that NUP76 is the functional orthologue to yeast Nup82 with a crucial role in mRNA export. In order to limit the possibility that the observed blockade of mRNA export is an indirect effect, i.e., the result of a disrupted pore architecture, we expressed a range of NUPs as N- or C-terminal eYFP fusions in the NUP76 auxin degron cell line to test whether their localisation to the pore is dependent on NUP76 (Figure 6 D and E and Figure S12). The localisation of the inner ring NUP96 was not affected by NUP76 depletion, suggesting that the overall pore structure remains intact (Figure 6D). Of the (putative) NUP76 associated proteins, only the pore localisation of NUP140 was clearly abrogated upon NUP76 depletion, while NUP149 and Gle2 still localised to the pore. Note that a slightly diminished pore localisation was observed for all four proteins, likely caused by the disrupted mRNA export and general loss in fitness rather than a specific impact on nuclear pore architecture. Thus, NUP140 localisation to the pore is fully dependent on NUP76, while NUP149 and Gle2 appear to be anchored independent of NUP76.

**Figure 6:**
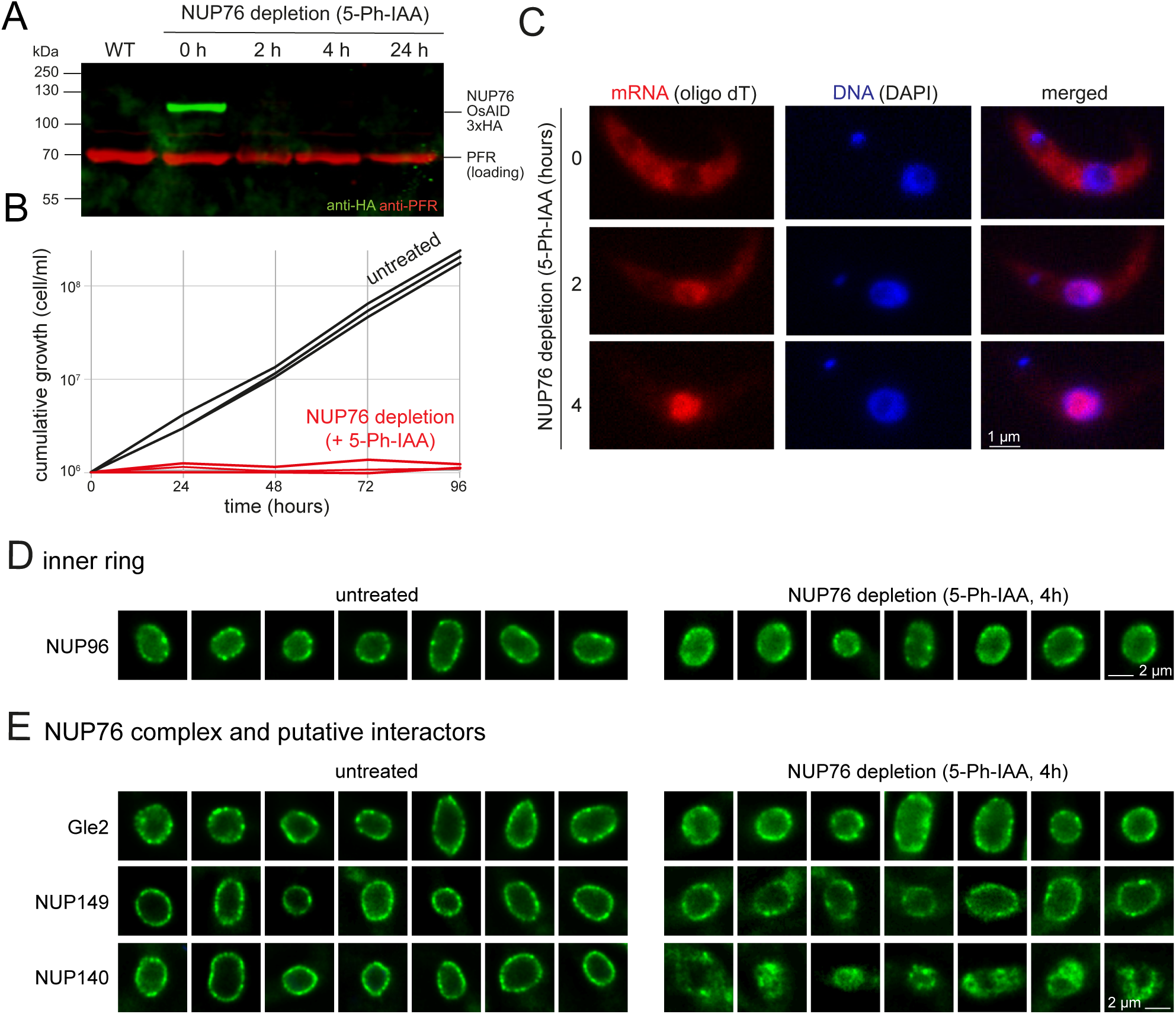
Depletion of TbNUP76 causes nuclear poly(A) accumulation and loss of NUP140 pore localisation. NUP76 protein was depleted using a degron system based on induction with the auxin derivative 5-Ph-IAA. Both alleles of NUP76 were replaced by NUP76 fused to OsAID-3HA at the C-terminus. **(A)** The depletion of the NUP76 protein at 2, 4, and 24 hours of induction was monitored on a western blot using anti-HA to detect NUP76 and anti-PFR as loading control. Wild type (WT) cells served as negative controls. Data of one representative clonal cell line are shown. **(B)** Growth was monitored over 4 days following NUP76 depletion. Data of three independent clonal cell lines are shown. **(C)** In situ hybridisation: cells were probed with oligo dT to monitor mRNA localisation. The DNA is labelled with DAPI. One representative cell is shown for untreated cells and cells after 2 and 4 hours of NUP76 depletion. **(D and E)** An N-terminal eYFP fusion of NUP96 and C-terminal eYFP fusions of NUP140, Nup149 and Gle2 were expressed from endogenous loci in the NUP76 degron cell line. The eYFP fluorescence of seven randomly selected nuclei is shown before and after induction of NUP76 depletion. Additional nuclei are shown in Figure S12.

In conclusion, the cytoplasmic-site localised NUP76 complex of trypanosomes, consisting of NUP76, NUP140 and NUP149, is distinct from the cytoplasmic mRNA remodelling complex of yeast and human. While NUP76 is the likely functional homologue to Nup82/NUP88 from yeast/human, NUP140 and NUP149 are trypanosome-unique with no sequence or structural homology to cytoplasmic site (filament) proteins from opisthokonts. The absence of a Nup159/NUP214 (yeast/human) orthologue in trypanosomes correlates with the absence of orthologues to its interaction partners Dbp5/DDX19 and Gle1/GLE1 (yeast/human), suggestive of significant differences of mRNA remodelling mechanisms in trypanosomes.

### A mechanistically divergent Ran-dependent mRNA export pathway in trypanosomes

Apart from the NUP76 complex, we mapped RANGAP and RANBP1 to the cytoplasmic site of the nuclear pore. Their sole cytoplasmic localisation is suggestive of a conserved function of these proteins in triggering GTP hydrolysis of RanGTP and thus disassembly of exportin - cargo-RanGTP and importin-RanGTP complexes. Trypanosome RanGAP is phylogenetically more closely related to a RabGAP (Gabernet-Castello *et al*, 2013) but its proposed function as RanGAP (Obado *et al*, 2016) is further corroborated by our study. Pore-anchoring of RanGAP and underlying mechanisms significantly vary across species, ranging from a SUMO dependent interaction with the metazoan specific RanBP2/NUP358 (Matunis *et al*, 1998; Mahajan *et al*, 1998, 1997; Matunis *et al*, 1996), over a WPP domain specific to plant RanGAP that interacts with a plant specific nucleoporin (Xu *et al*, 2007; Rose & Meier, 2001), to no pore localisation at all in *S. cerevisiae* and *S. pombe* (Hopper *et al*, 1990; Melchior *et al*, 1993). The mechanism of pore localisation of trypanosome RanGAP is thus likely unique and may involve interactions with likewise unique trypanosome specific NUPs such as NUP140 and/or NUP149. Trypanosome RanBP1 consists of a disordered 30-amino acid stretch followed by a conserved RanBP domain (Figure S13A) and it remains unclear whether it has binding sites to the pore or simply concentrates in sites of cargo docking.

The lack of a Dbp5 homologue and the interaction between Mex67 and Ran implies that trypanosomes employ the Ran-GTP gradient for mRNA export (Obado *et al*, 2016). In fact, while Ran is predominantly nuclear localised in humans (Hutchins *et al*, 2009) we find biotinylated Ran targets on both sites of the pore, possibly reflecting Ran engagement in cargo import and export (Hutchins *et al*, 2009). This unique dual usage of the Ran pathway for both mRNA and protein cargo presents a formidable challenge for export/import ratio moderation. It is tempting to speculate that RanGTP is anchored at the basket site awaiting the MEX67 bound mRNP, which is then liberated on the cytoplasmic site driven by RanGAP-catalysed GTP hydrolysis. The lack of Dbp5 suggests that ATP-dependent mRNP disassembly at the cytoplasmic site of the pore is dispensable in trypanosomes implying a fundamentally different mode of interaction between MEX67 and mRNA. Indeed, trypanosome Mex67 uniquely carries a CCCH-type zinc finger instead of the canonical RNA recognition motif containing RNA binding domain (RRM/RBD) found in ophistokonts (Kramer *et al*, 2010; Dostalova *et al*, 2013), and trypanosomes lack mRNA adaptors (SR proteins) that would require stripping during cytoplasmic remodelling. Thus, the exported trypanosome RNP may exhibit lower stability and complexity, making a remodelling RNA helicase redundant.

While only five proteins are specific to the cytoplasmic site, the nuclear site of the pore appears more complex. Next to the previously described basket proteins NUP110 and NUP92 (Obado *et al*, 2016; DeGrasse *et al*, 2009) we found nuclear site localisation for the FG nucleoporins NUP64 and NUP98, as well as for RanBPL, putative TREX-complex proteins and several further proteins that are mostly unique to trypanosomes.

The FG-NUPs NUP64 and NUP98 are unique to trypanosomes and in a complex with NUP75 (Obado *et al*, 2016) which appears to extend to the inner ring. NUP64 and NUP98 were previously suggested to be the (functional) orthologues of *S. cerevisiae* Nup1 and Nup60, as they carry the same FG-type and engage in an interaction with the putative Sac3 homologue (Obado *et al*, 2016; Butterfield *et al*, 2024). Our data now show the exclusive nuclear-side position of these NUPs, in full support of this model.

RanBPL has a Ran-binding domain which is very similar to the one of cytoplasmic-site localised RanBP1 but has a longer disordered amino-terminal stretch (Figure S13) and was previously characterized as Ran binding protein with a clear preference to RanGTP over RanGDP (Brasseur *et al*, 2014). Thus, RanBPL may be the trypanosome functional counterpart to basket proteins Nup2/NUP50 (yeast/human), which also possess Ran-binding domains. Whilst the multiple roles of Nup2/NUP50 remain largely elusive, one known function is the acceleration of protein import complex disassembly through stimulation of RanGEF/RCC1 activity (Holzer & Antonin, 2022), analogous to the function of RanBP1 as enhancer of RanGAP activity at the cytoplasmic site (Seewald *et al*, 2003). In trypanosomes, a RanGEF has not yet been identified and the absence of a detectable RanGEF/RCC1 domain among the proteins biotinylated by Ran indicates that a trypanosome RanGEF/RCC1 is either absent or divergent. Theoretically, RanBPL has the potential to compensate for the absence of the canonical RanGEF/RCC1: instead of directly catalysing the GDP to GTP exchange, RanBPL1 could act by stabilizing RanGTP and preventing GTP hydrolysis, driving the equilibrium towards RanGTP. However, the observation that RanBPL silencing evokes only a mild growth phenotype (Brasseur *et al*, 2014) argues against this hypothesis. Altogether, our study fortifies the hypothesis of a Ran-dependent mRNA export pathway in trypanosomes and opens new avenues for exploration of the underlying molecular mechanisms.

## CONCLUSION

Our revisited map of the trypanosome nuclear pore conforms to the pattern of conservation at the core scaffold regions and diversity at the borders of the pore (Makarov *et al*, 2021). We discovered an asymmetric architecture, confidently placing the NUP76 complex exclusively to the cytoplasmic site and defining the sole localisation of the trypanosomatid-exclusive FG NUPs NUP64 and NUP98, at the basket site. Notably, this corrects the current view of a largely symmetric trypanosome nuclear pore and ultimately supports moderation of directional nucleocytoplasmic transport which is crucially dependent on asymmetric components at the nuclear pore borders in other systems. For the NUP76 complex, our data strongly indicates a crucial function as cytoplasmic mRNP remodelling hub, analogous to the Nup82/NUP88 complex in opisthokonts, while the presence of trypanosome-unique NUP140 and NUP149 implies significant mechanistic difference. Mapping of the export factors Mex67 and Ran elucidated further divergence, supporting a trypanosome-specific, Ran-dependent export system. Lastly, we present a comprehensive assignment of pore localised proteins to sub-regions of the nuclear pore that resulted in the identification of novel nuclear pore components, including three putative members of a trypanosome TREX-2 complex. Altogether, our approach delivers asymmetric and novel nuclear pore components, including positional information, which can now be interrogated for functional roles to explore trypanosome-specific adaptions of nuclear transport, export control, and mRNP remodelling.

The compartmentalisation of hereditary information in the nucleus necessitated the invention of a gateway allowing mRNPs and a variety of other essential cargos to cross the nuclear envelope. Hence, the nucleus, eponymous feature of eukaryotes, must have co-evolved with the nuclear pore that constitutes this gateway. The study of nuclear pores in evolutionary divergent eukaryotes, such as the ancient trypanosomes, is fundamental to understand the evolutionary origin of the nucleus and decipher the complexity of the nuclear pore as platform with multiple cargo routes. Our study contributes a roadmap of the trypanosome nuclear pore, reporting conserved and non-conserved features and devising a plethora of new leads for further exploration.

## SUPPLEMENTAL FIGURES

**Figure S1:** Establishment and validation of the expansion microscopy protocols.

**Figure S2:** Additional proExM images of the NUP76 complex proteins.

**Figure S3.** UExM images of NUP64-4Ty1 with NUP76 complex proteins tagged with 3xHA (labelled using antibodies).

**Figure S4:** Statistical analysis of TurboID experiments

**Figure S5:** Inducible overexpression of Mex67 (growth curves and Western blotting to check for expression levels).

**Figure S6:** Nucleolus can be identified by reduction in DAPI stain.

**Figure S7:** Mapping proteins with the proximity map: Karyopherins.

**Figure S8:** Mapping proteins with the proximity map: unlabelled proteins.

**Figure S9:** Structures of yeast NUP82, human NUP88 and predicted *T. brucei* NUP76.

**Figure S10:** The NUP149 zinc fingers in comparison to the ones from human NUP153 and NUP358.

**Figure S11:** Verification of NUP76 auxin degron cell lines by diagnostic PCR.

**Figure S12:** Effect of NUP76 depletion on NUP140, NUP149, Gle2 and NUP96 localisation: additional images.

**Figure S13:** AlphaFold2 models of RanBP1 and RanBPL

## ACKNOWLEDGEMENTS

The project was funded by a bilateral GACR/DFG grant (project IDs.: 21-19503J and KR4017/9-1; to M. Z. and S. K., respectively) and the DFG grant KR4017/8-1 to S.K. We are grateful to the OMICS Proteomics BIOCEV core facility for excellent technical service. We like to thank Eva Kowalinski (EMBL, Grenoble, France) for expert help with Alphafold and Colabfold. We are grateful to Mark Carrington for providing the highly useful auxin degron system.

