## Supplementary material for "A combination of expansion microscopy and proximity labelling reveals conserved and unique asymmetric functional hubs at the trypanosome nuclear pore": Sup Figures

### Figure S1 (page 1/2)

#### A principle of ProExM

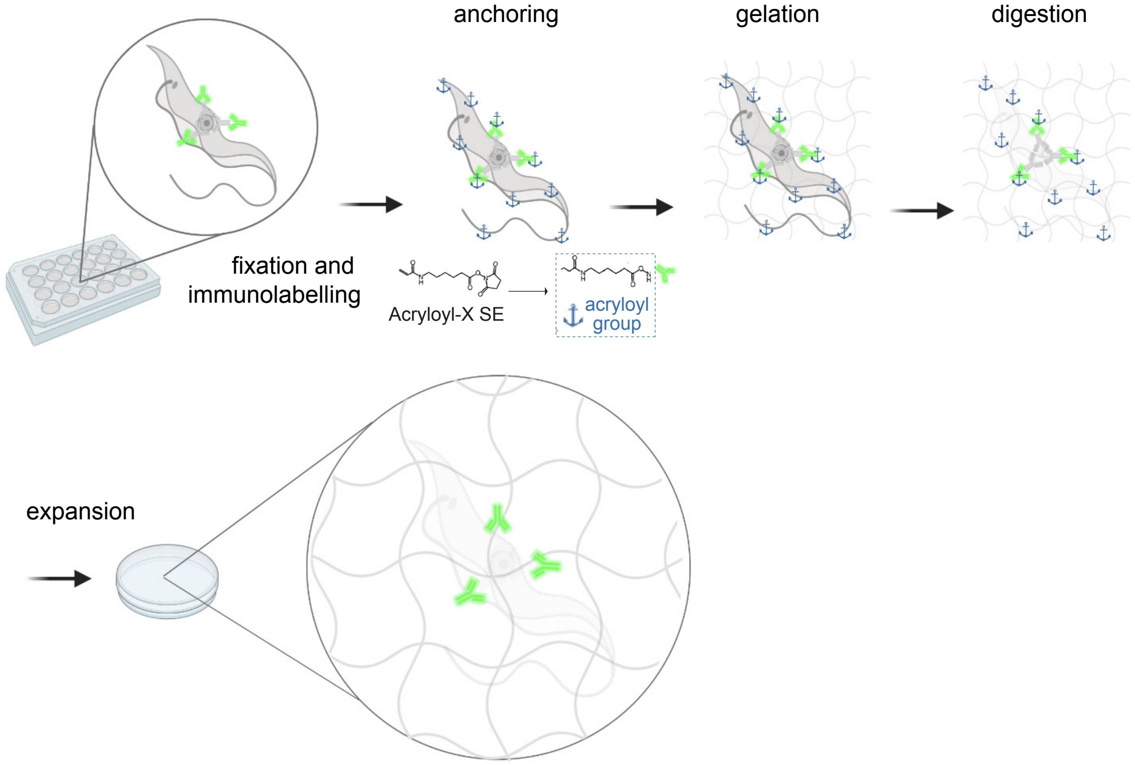


#### B principle of UExM

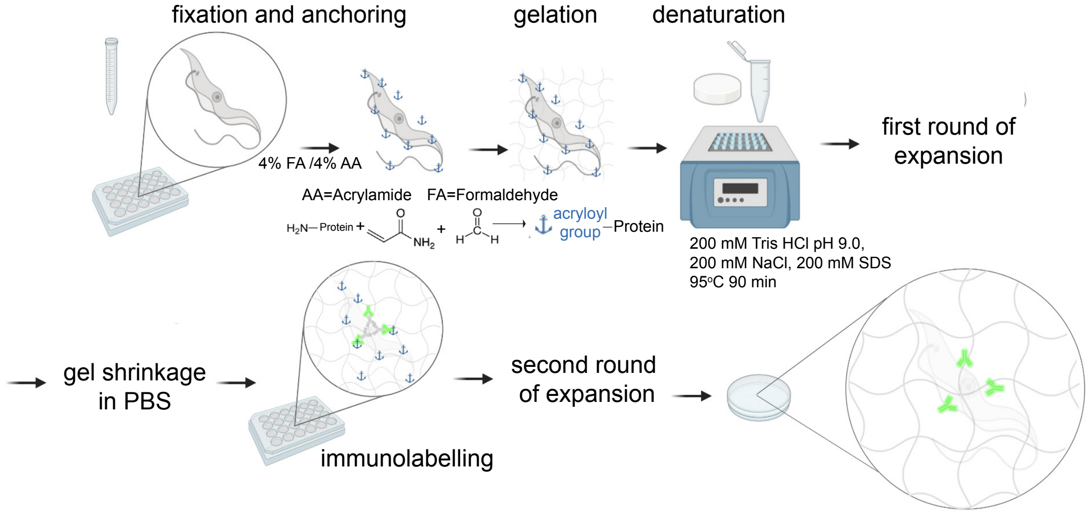


#### C Expansion factor

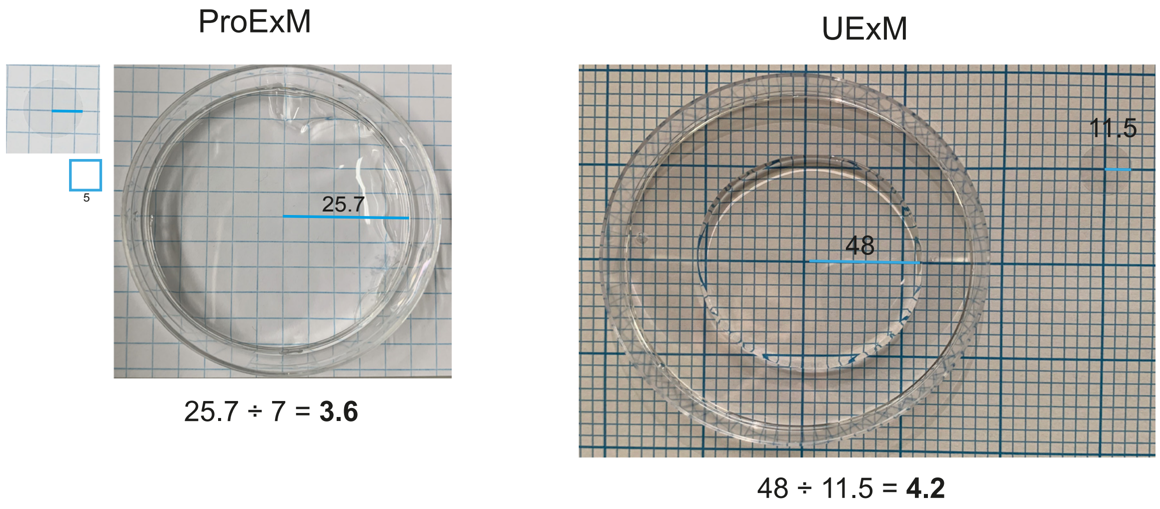

D Test for isotropic expansion

| | no expansion | proExM | UExM |
|---|---|---|---|
| example image: | | | |

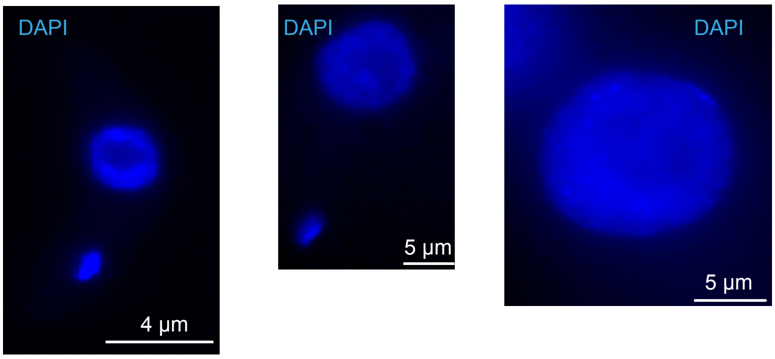


| | no expansion | proExM | UExM |
|---|---|---|---|
| diameter of nuclei ± SD: | 2.4 ± 0.2 µm (n = 21) | 8.4 ± 0.7 µm (n =72) | 11.5 ± 1.1 µm (n = 52) |
| expansion factor based on nuclei: | | 8.4 ÷ 2.5 = **3.5** | 11.5 ÷ 2.5 = **4.6** |

**Figure S1: Validation of Protein Retention Expansion Microscopy (proExM) and Ultrastructural Expansion Microscopy (UExM) (A)** Workflow of the proExM method, adapted from[1]. **(B)** Workflow of the UExM method, adapted from[2]. **(C)** Calculation of the expansion factor for proExM (left) and UExM (right) by comparing the size of the non-expanded and the expanded gel, using squared paper as a size reference. Measurements were done with Fiji[3]. **(D)** The diameter of DAPI stained nuclei was measured in unexpanded cells and after expansion. The so-measured expansion factor is very similar to the one measured by the increase in gel size (C), indicating isotropic expansion of the nucleus.

[1]Asano SM, Gao R, Wassie AT, Tillberg PW, Chen F & Boyden ES (2018) Expansion Microscopy: Protocols for Imaging Proteins and RNA in Cells and Tissues. Curr Protoc Cell Biol 80
[2]Gambarotto D, Hamel V & Guichard P (2021) Ultrastructure expansion microscopy (U-ExM). In pp 57–81.
[3]Schindelin J, Arganda-Carreras I, Frise E, Kaynig V, Longair M, Pietzsch T, Preibisch S, Rueden C, Saalfeld S, Schmid B, et al (2012) Fiji: an open-source platform for biological-image analysis. Nat Methods 9: 676–682

Figure S2

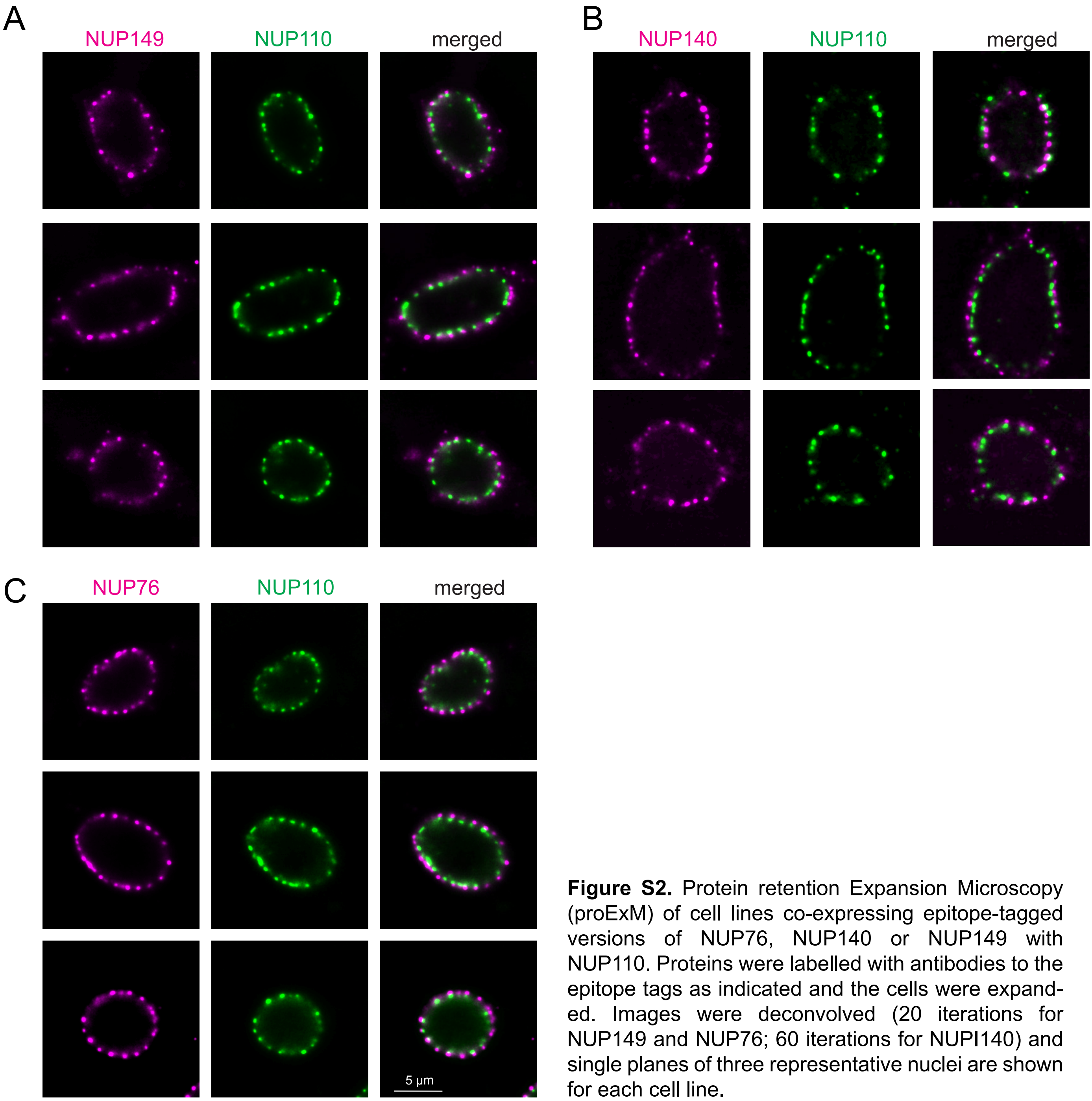


**Figure S2.** Protein retention Expansion Microscopy (proExM) of cell lines co-expressing epitope-tagged versions of NUP76, NUP140 or NUP149 with NUP110. Proteins were labelled with antibodies to the epitope tags as indicated and the cells were expanded. Images were deconvolved (20 iterations for NUP149 and NUP76; 60 iterations for NUPI140) and single planes of three representative nuclei are shown for each cell line.

Figure S3

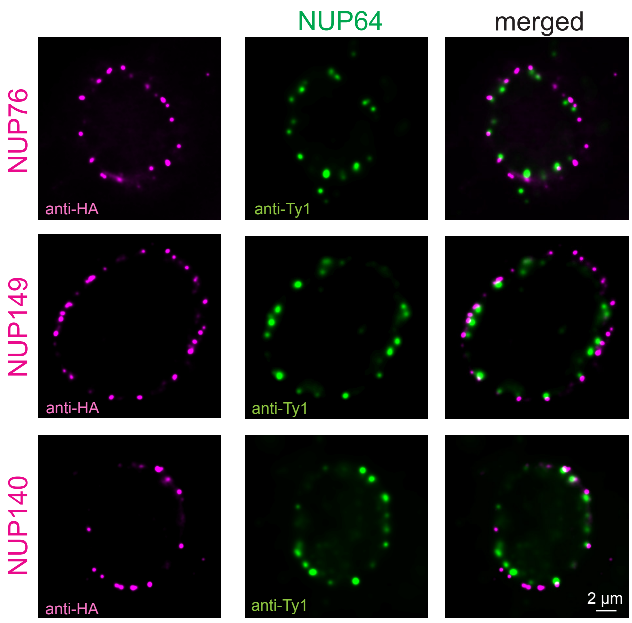


**Figure S3.** Ultrastructure Expansion Microscopy (UExM) of lines co-expressing HA-tagged versions of NUP76/NUP140/NUP149 with NUP64-4Ty1. Labelling was done with anti-Ty1 and anti-HA. Images were deconvolved with 60 iterations and a single plane image of one nucleus of each cell line is shown.

### Figure S4

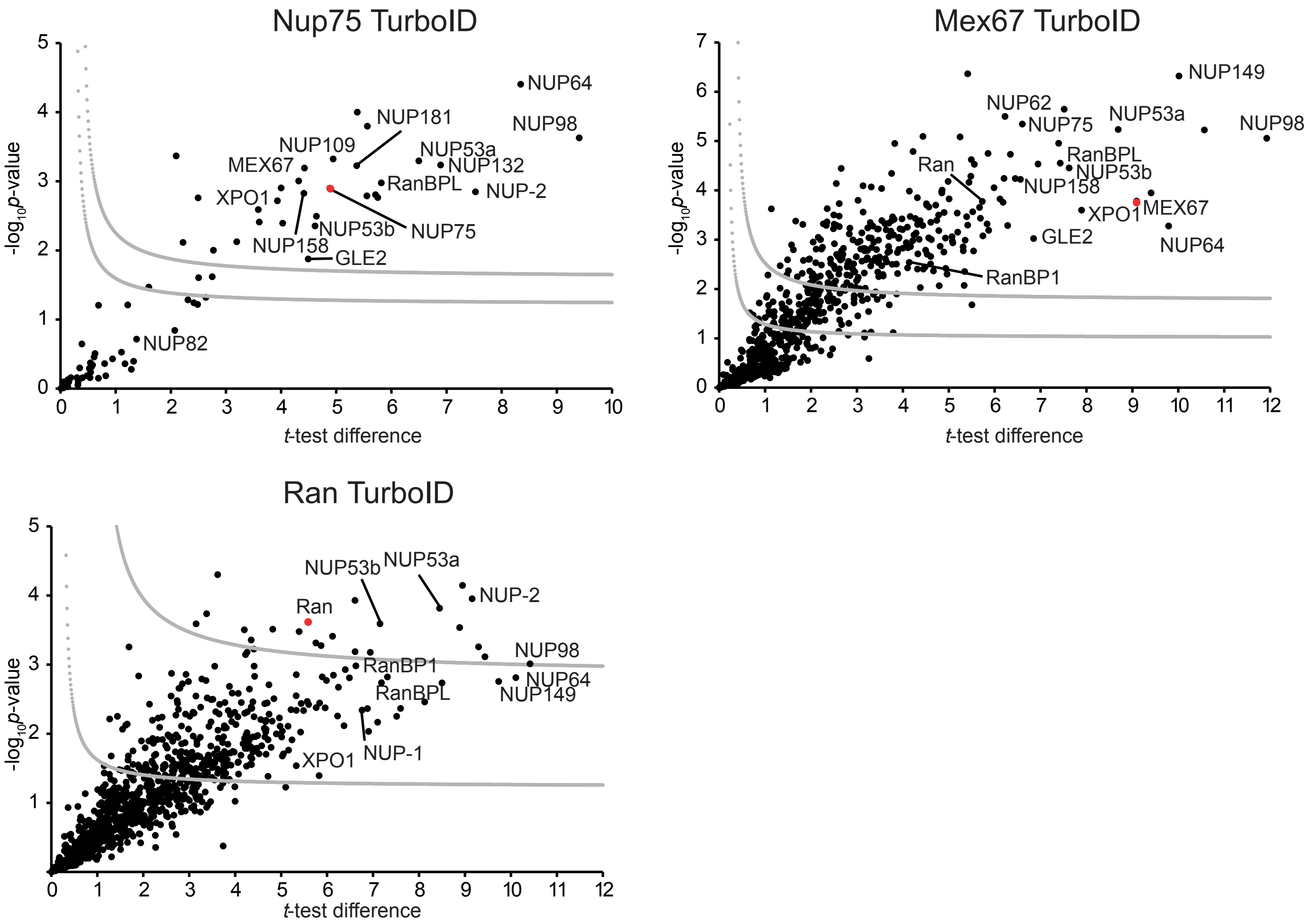


**Figure S4: Statistical analysis of TurboID experiments.**
Hawaii plot (multiple volcano plots) of label-free quantification results of the BioID experiments for NUP75, Ran and Mex67 with fused TurboID tag at the C-terminus. All samples were prepared at least in duplicate. To generate the volcano plots, the $-\log_{10}p$-value was plotted versus the t-test difference (difference between means), comparing each respective bait experiment to the wt control. Potential interactors were classified according to their position in the plot, applying cut-off curves for "significant class A" (SigA; gray, upper curve; FDR = 0.01, s0 = 0.1) and "significant class B" (SigB; gray, lower curve; FDR = 0.05, s0 = 0.1), respectively. Bait proteins are indicated by a red dot, selected known NUPs and transport factors are labelled and LFQ data is given in Table S2.

Figure S5

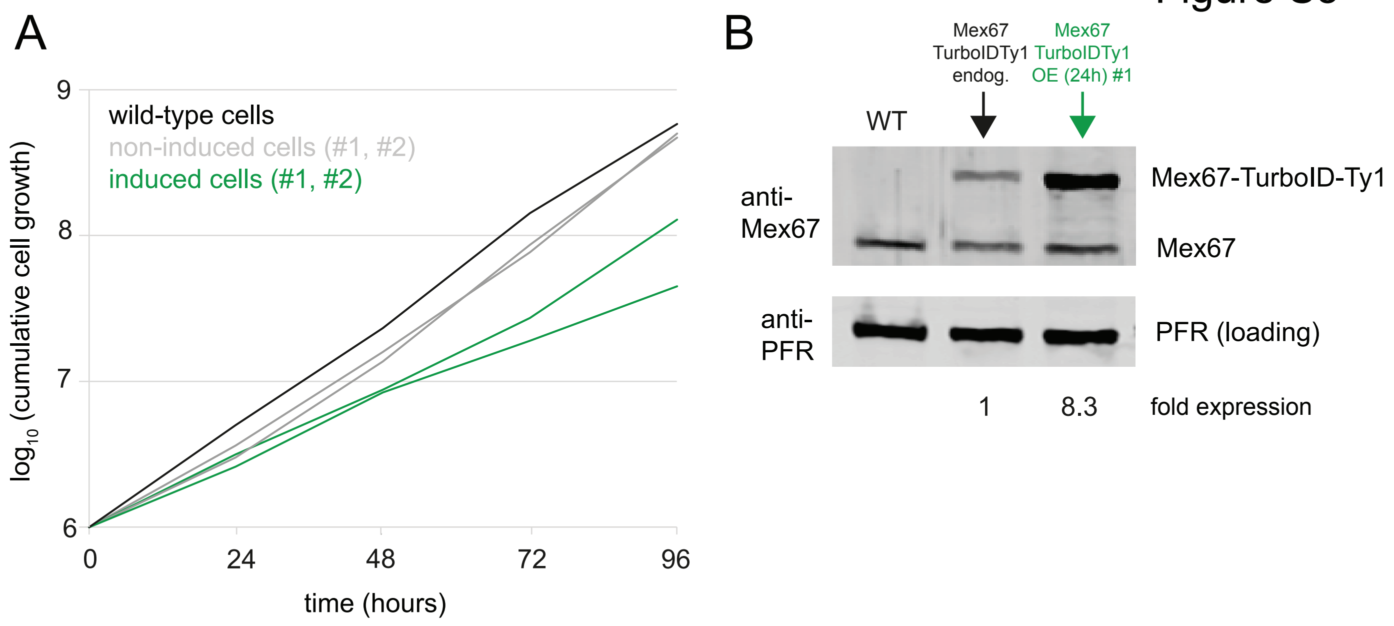


**Figure S5. Inducible overexpression of Mex67-TurboID-Ty1**
(A) Growth of wild-type cells and of two clones with the inducible expression of Mex67-TurboID-Ty1 from an ectopic locus without (gray) and with (green) induction using tetracyclin. Growth was monitored for 96 hours with daily measurements. (B) Western blot loaded with cell lysates of wild-type cells, cells expressing Mex67-TurboID-Ty1 from the endogenous locus and cells with induced expression of Mex67-TurboID-Ty1 from an ectopic locus for 24 hours (clone #1). The Western blot was probed with antibodies specific to Mex67 and to PFR (to control for loading). The intensities of the bands were quantified using hte LiCor Odyssey software and the extent of overexpression was quantified, normalised by the loading control.

### Figure S6

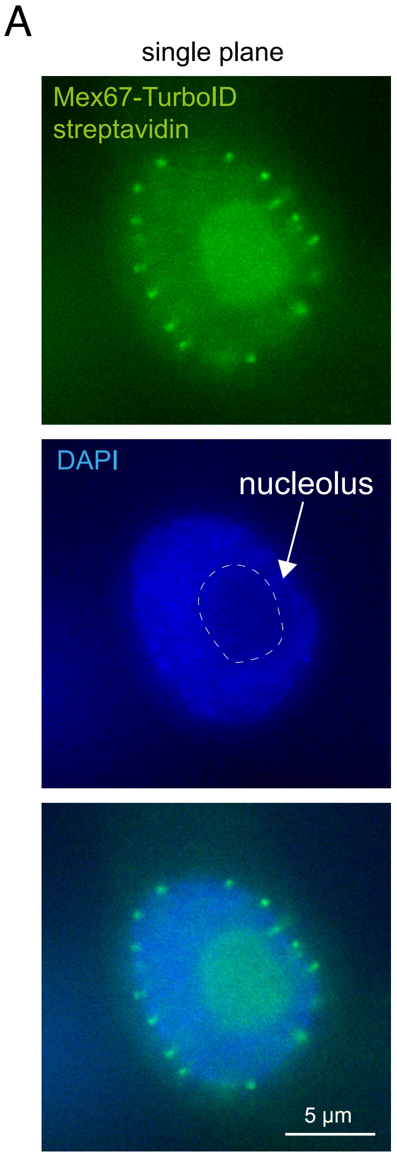


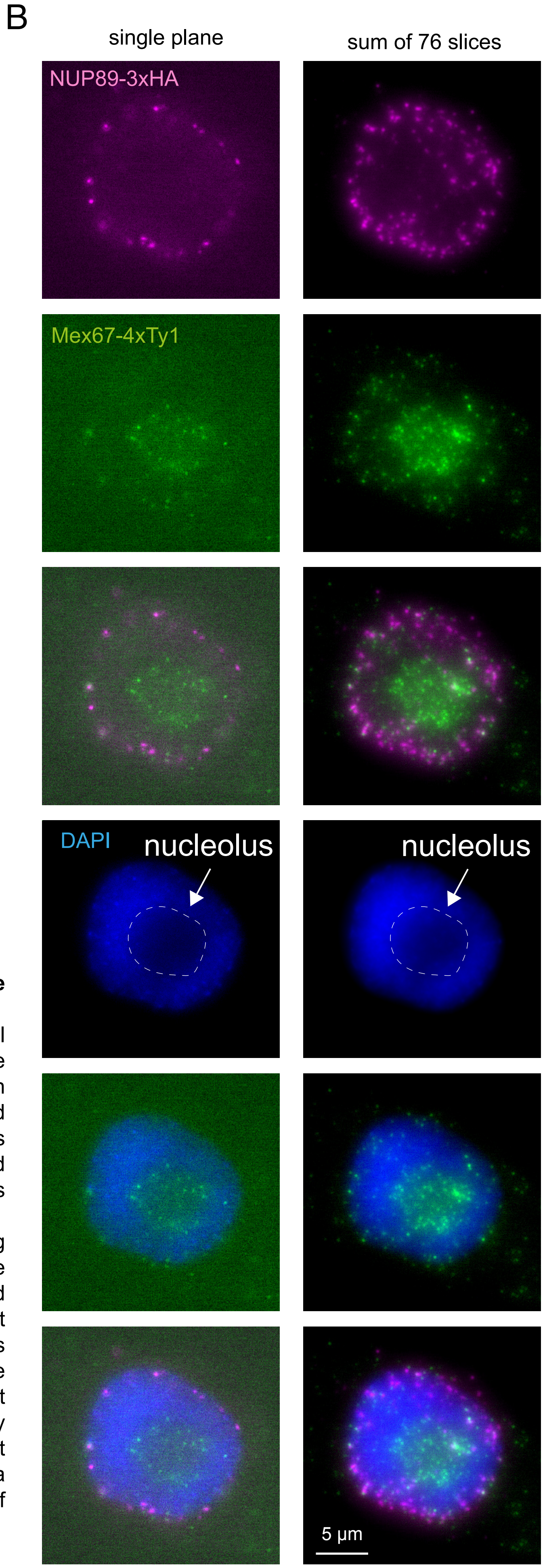


**Figure S6. Mex67 does localise to the nucleolus.**
(A) Expansion microscopy image (UExM) of a cell overexpressing Mex67 fused to TurbID (compare to Figure 3B). Mex67 biotinylation is imaged with fluorescent streptavidin and the DNA is stained with DAPI. The single trypanosome nucleolus is defined by the reduction in DAPI signal and colocalises with Mex67. A single plane image is shown as raw data (no deconvolution).
(B) UExM image of a cell co-expressing NUP89-3HA and Mex67-4Ty1. The proteins were detected with antibodies (anti-HA; anti-Ty1) and the DNA is stained with DAPI. Mex67, but not NUP89 has signal at the nucleolus. This pvoves that the nucleolar streptavidin signal of the Mex67-TurboID fusion reflects the correct localisation of Mex67 and is not caused by biotinylation of Mex67 interactors, at least not entirely. A raw (non-deconvolved) image of a single plane is shown, as well as a projection of 76 slices from a Z-stack (sum slices).

### Figure S7 page 1/2

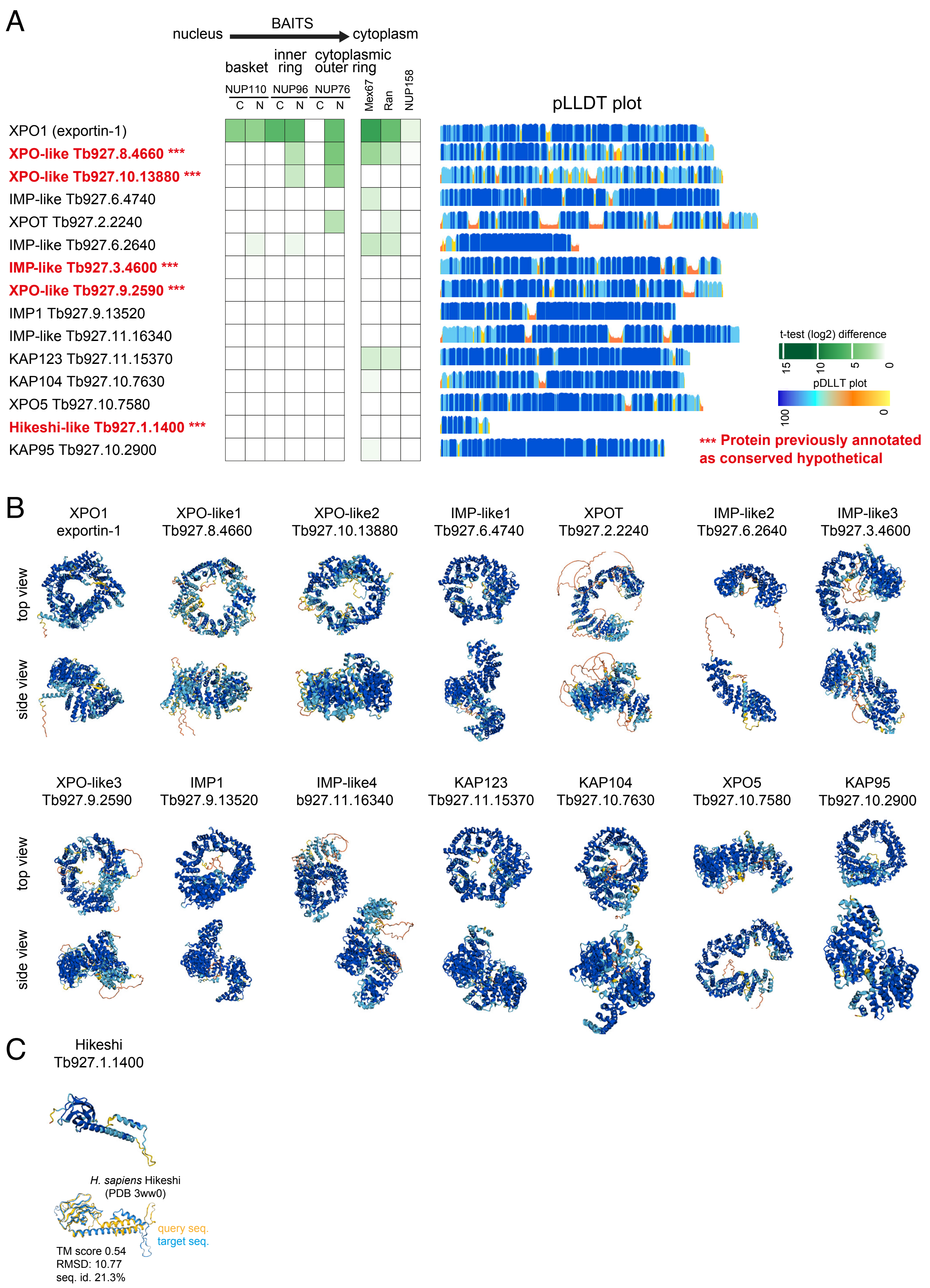

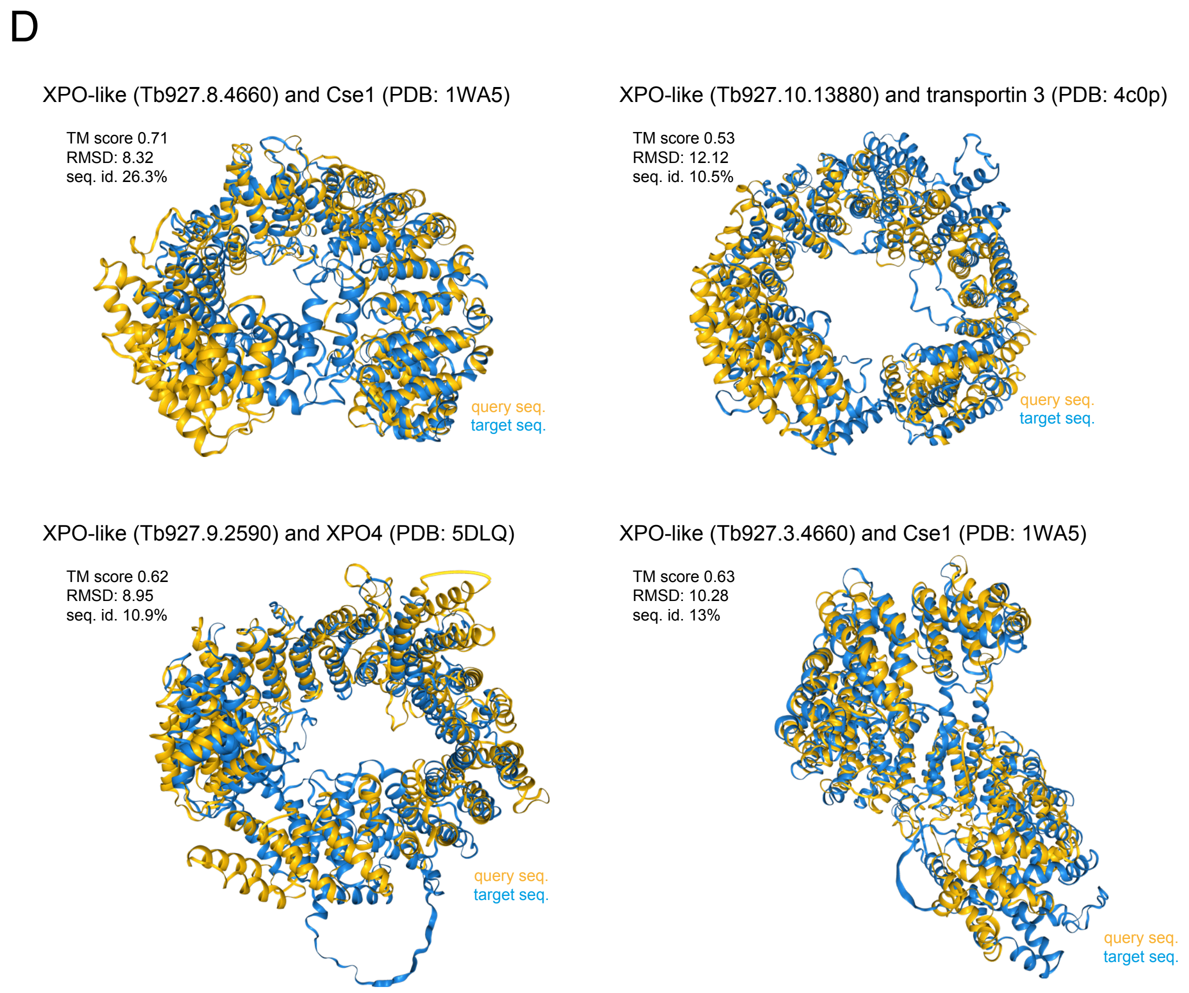


**Figure S7: Karyopherins are poorly labelled by NUP110, NUP96 and NUP76**
(A) All trypanosome proteins with nuclear pore localisation predicted by TrypTag[1] were screened for the presence of importin/exportin folds by running their AlphaFold2 models[2] in FoldSeek[3]. This resulted in the identification of 15 importins/exportins, of which five were not previously annoted as such (***** in A**). Mass spectrometry data from proximity labelling experiments of NUP110, NUP96, NUP76, NUP158, Ran and Mex67 were analysed and the labelling of these karyopherins is shown with a color-code (t-test difference values, log2-transformed). pLLDT plots are included.

(B) Trypanosmatid-optimised AlphaFold2 models of 14 karyopherins (top and side view of the models, taken from[2] and coloured based on pLLDT values shown in (A).

(C) Trypanosmatid-optimised AlphaFold2 models of Hikeshi (taken from[2] and coloured based on pLLDT values shown in (A) and the FoldSeek outputs[3]: superimposition between the AlphaFold2 models of the trypanosomatid orthologues (target sequence, coloured blue) and the FoldSeek best hit on the PDB database (query sequence, coloured yellow), with root mean square deviation of atomic positions (RMSD) of the superimposition, internal confidence values (template modelling scores, TM, ranging from 0-1, from worst to best) and sequence identity values below.

(D) The four proteins with structural homologies to karyopherins that were not annotated as karyopherins (***** in A**) and their FoldSeek search outputs[3]: superimposition between the AlphaFold2 models of the trypanosomatid orthologues (target sequence, coloured blue) and the FoldSeek best hit on the PDB database (query sequence, coloured yellow), with root mean square deviation of atomic positions (RMSD) of the superimposition, internal confidence values (template modelling scores, TM, ranging from 0-1, from worst to best) and sequence identity values below.

[1] Billington, K. et al. Genome-wide subcellular protein map for the flagellate parasite Trypanosoma brucei. Nat. Microbiol. 8, 533–547 (2023).
[2] Wheeler R. A resource for improved predictions of Trypanosoma and Leishmania protein three-dimensional structure. PLoS ONE 16(11): e0259871 (2021).
[3] van Kempen, M. et al. Fast and accurate protein structure search with Foldseek. Nat Biotechnol. 42, 243–246 (2024).

Figure S8

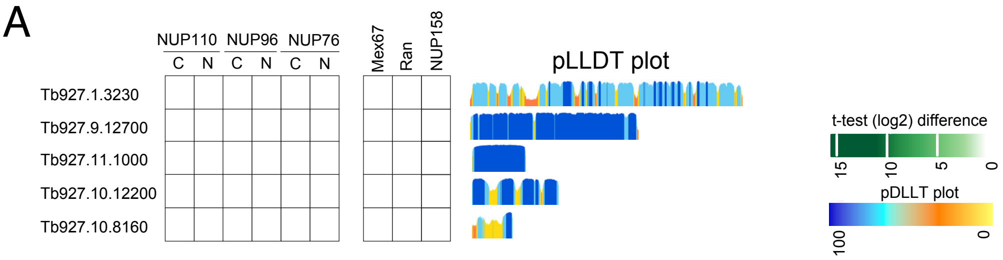


B These images are from the Tryptag genome-wide localisation database.

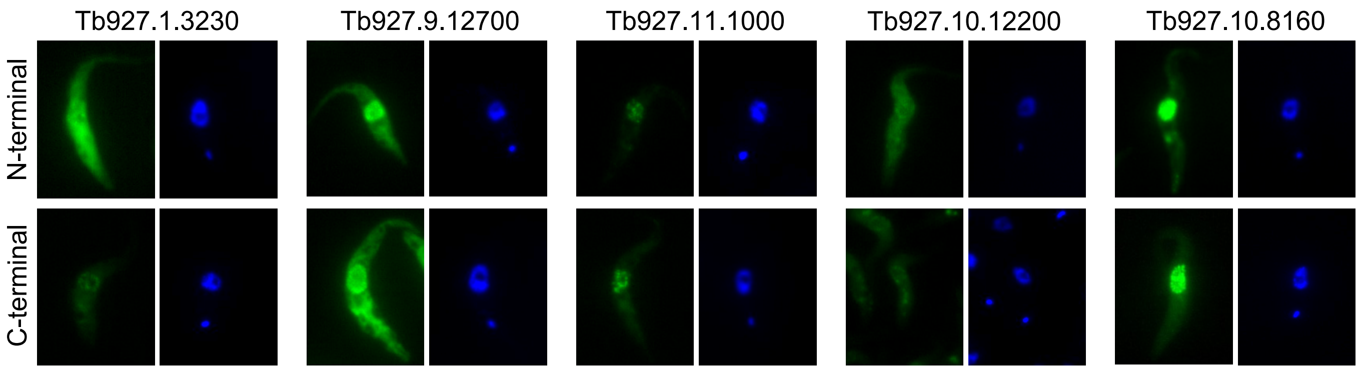


**Figure S8: Proteins with nuclear pore localisation that are not labelled by NUP110/96/110, Mex67, Ran or NUP158**
**(A)** Mass spectrometry data from proximity labelling experiments of NUP110, NUP96, NUP76, NUP158, Ran and Mex67 were analysed for proteins with nuclear pore localisation[a]. Five proteins were not labelled by either of these six proteins, and the pLLDT plots are shown.
**(B)** Localisation images of Neon-green fusions of these proteins were taken from TrypTag[a]. The DNA is stained with DAPI and shown in blue. Images of both N-terminal and C-terminal Nean-green fusions are shown.

[a] Billington, K. et al. Genome-wide subcellular protein map for the flagellate parasite Trypanosoma brucei. Nat. Microbiol. 8, 533–547 (2023).

### Figure S9

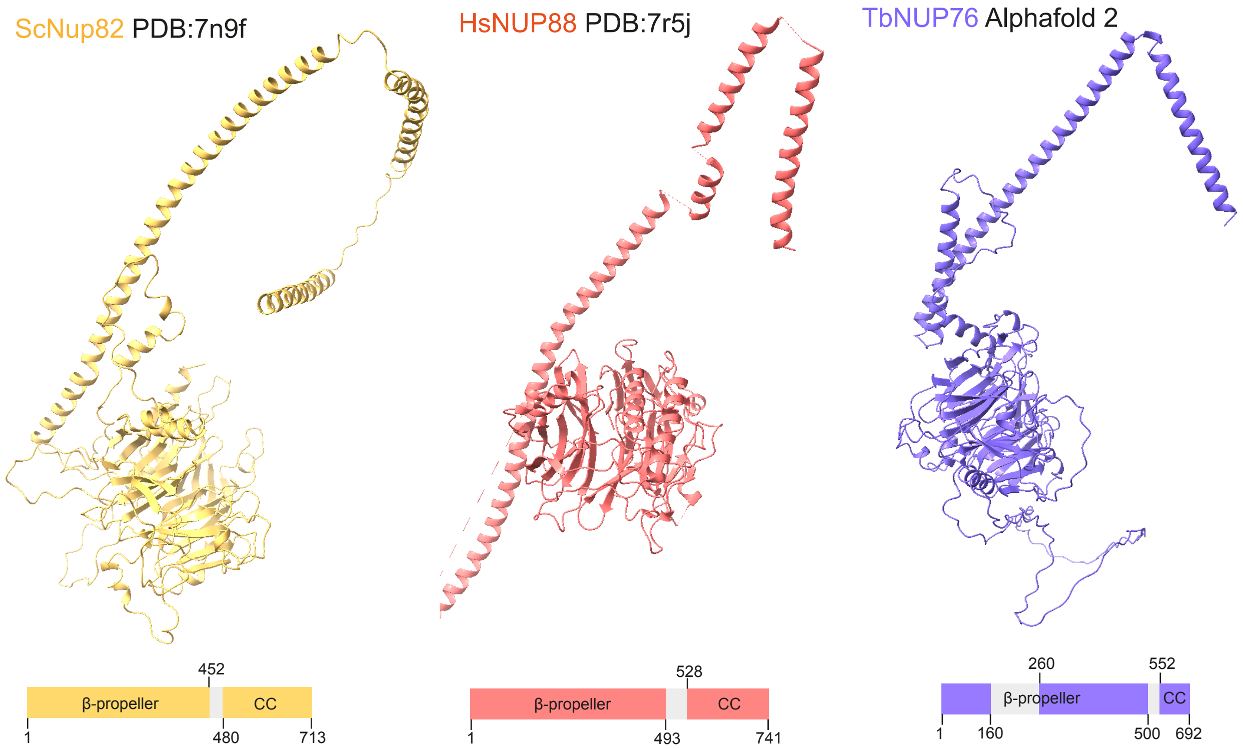


**Figure S9:** Homologies between experimentally resolved structures of ScNup82[1], HsNUP88[2] and the AlphaFold2-predicted model of TbNUP76[3]. All three proteins have very similar structures, namely an N-terminal beta-propeller that is followed by coiled-coil (CC) domains. One difference is the long disordered region within the beta-propeller of TbNUP76, that is not present in *Sc*NUP82 and HsNUP88. Below each structure is a schematic drawing.

1 Akey CW, Singh D, Ouch C, Echeverria I, Nudelman I, Varberg JM, Yu Z, Fang F, Shi Y, Wang J, et al (2022) Comprehensive structure and functional adaptations of the yeast nuclear pore complex. Cell 185: 361-378.e25

2 Mosalaganti S, Obarska-Kosinska A, Siggel M, Taniguchi R, Turoňová B, Zimmerli CE, Buczak K, Schmidt FH, Margiotta E, Mackmull M-T, et al (2022) AI-based structure prediction empowers integrative structural analysis of human nuclear pores. Science (1979) 376

3 Wheeler RJ (2021) A resource for improved predictions of Trypanosoma and Leishmania protein three-dimensional structure. PLoS One 16: e0259871

### Figure S10

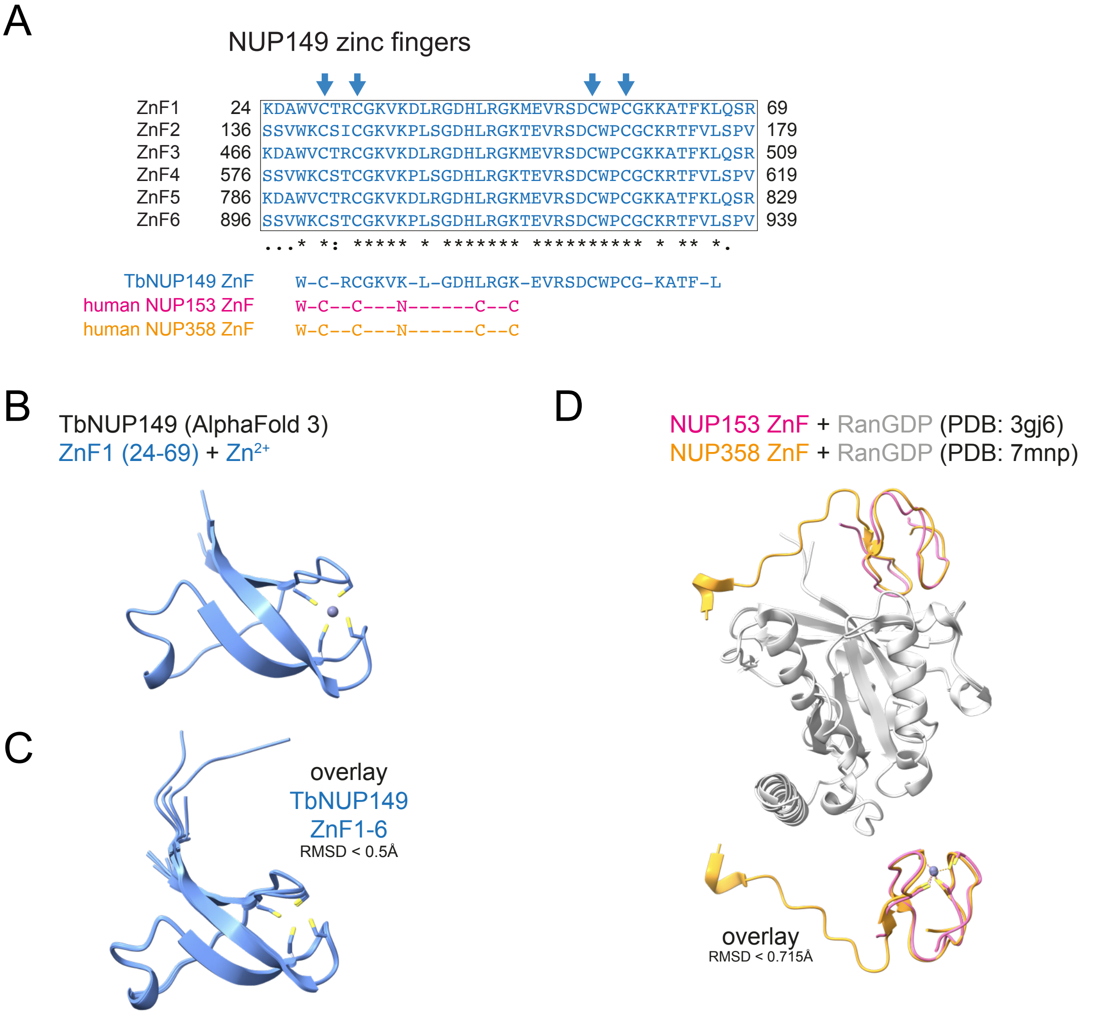


**Figure S10:**

**(A)** Alignment of the six zinc fingers of *T. brucei* NUP149 with consensus sequence. The consensus sequence of the unrelated zinc fingers of the human NUP153 and NUP358 (taken from [1]) is shown for comparison. Arrows point to the cysteine residues used to chelate the $Zn^{2+}$ ion (shown in B).

**(B)** AlphaFold3[2] model of the most N-terminal zinc finger of NUP149 (Zn1, amino acids 24-69) together with a zinc ion. The cysteine side chains are shown in yellow.

**(C)** Trypanosome-optimised AlphaFold2 models of all six zinc fingers of NUP149 were superimposed, using the AlphaFold2 structure of the first zinc finger as a reference structure (RMSD of all superimpositions are all below 0.5Å). All cysteine side chains are shown in yellow.

**(D)** Overlay of the experimentally resolved structures of the zinc fingers of human NUP153[3] and human NUP358[4] in complex with RanGDP. Note that there is no similarity to the zinc fingers of *T. brucei* NUP149 (compare B and C).

[1] Partridge, J. R. & Schwartz, T. U. Crystallographic and Biochemical Analysis of the Ran-binding Zinc Finger Domain. J. Mol. Biol. 391, 375–389 (2009).
[2] Abramson, J. et al. Accurate structure prediction of biomolecular interactions with AlphaFold 3. Nature 630, 493–500 (2024).
[3] Partridge, J. R. & Schwartz, T. U. Crystallographic and Biochemical Analysis of the Ran-binding Zinc Finger Domain. J. Mol. Biol. 391, 375–389 (2009).
[4] Bley, C. J. et al. Architecture of the cytoplasmic face of the nuclear pore. Science 376, eabm9129 (2022).

### Figure S11

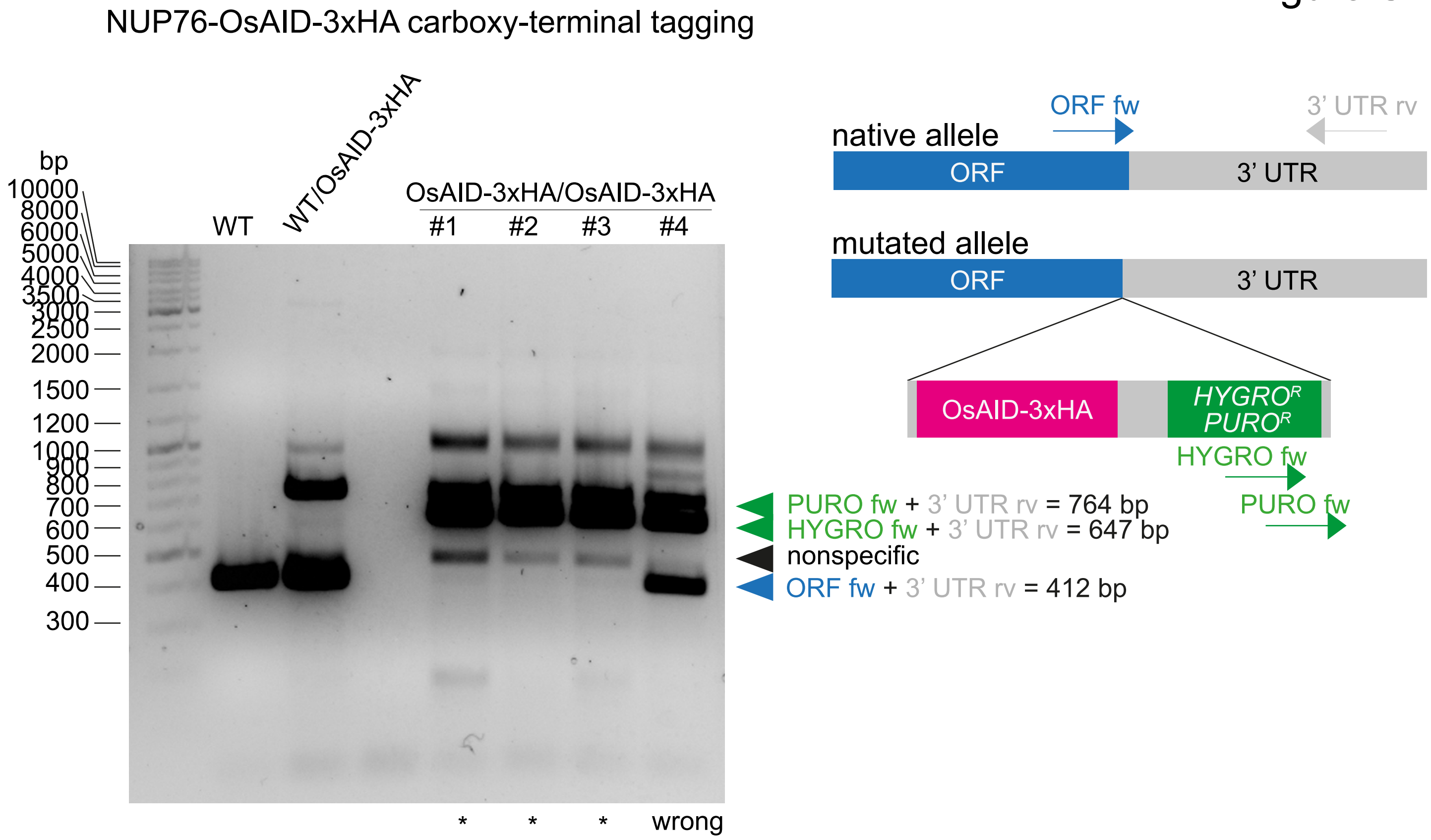


**Figure S11: Confirmation of NUP76-OsAID-3xHA homozygous cell lines by diagnostic PCR**
(A) The auxin inducible degron system was employed for inducible degradation of *T. brucei* NUP76. Both endogenous alleles of the NUP76 gene were fused to OsAID-3xHA at the carboxy-terminus. Two resistance cassettes were used, one with a puromycin and the other with a hygromycin resistance gene. The PCR strategy that was used to evaluate the cell line and, in particular, to control for the absence of the wild type allele, is schematically pictured (right). PCR reactions were performed with a mixture of all three forwards oligos and the reverse oligo and products are resolved on an agarose gel. A non-specific band is indicated with a black triangle. The wild type and the mutated bands are indicated with a blue and green triangles, respectively. Three clones are positive (#1-3, labelled with *) and (#4) is wrong.

Figure S12

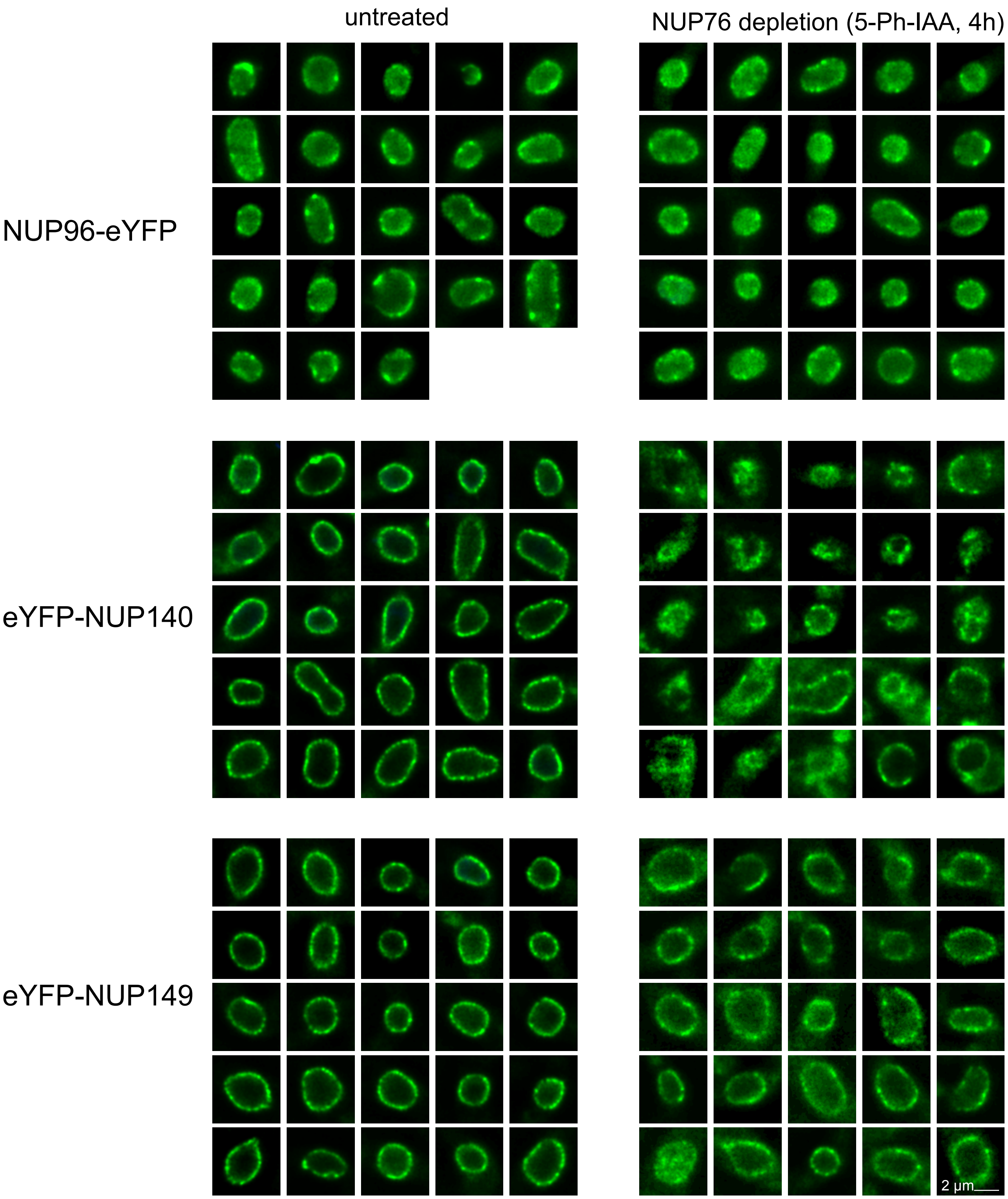


**Figure S12:** Additional images for Figure 6D and E.

### Figure S13

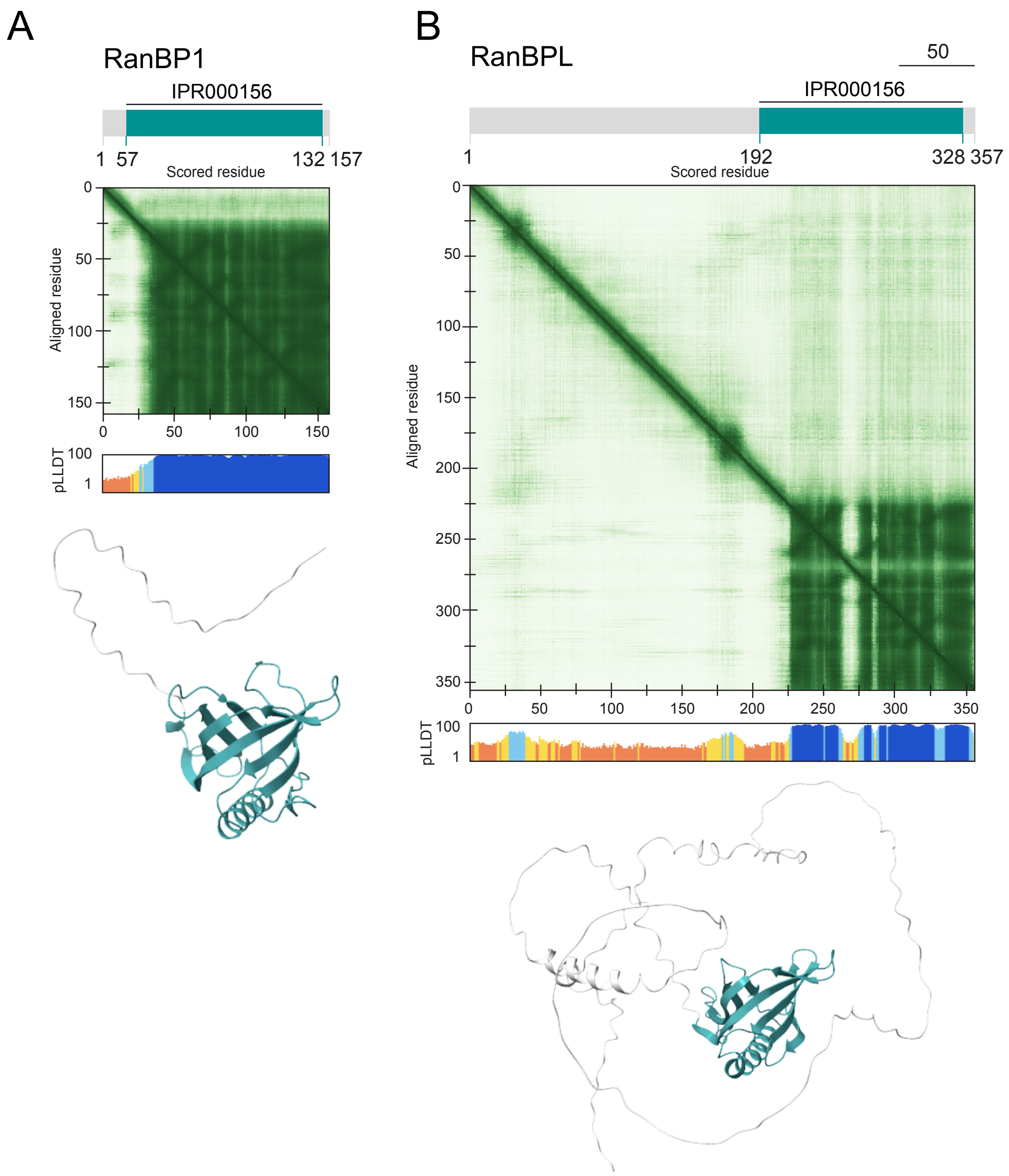


**Figure S13: Trypanosome RANBP1 and RANBPL**
Models of trypanosome-optimised AlphaFold2 predictions of RANBP1 **(A)** and RANBPL **(B)** pAE plots, PLLDT plots and the predicted structures are shown, with structured parts coloured and disordered regions shown in grey.
